# A potential role for acyl-phosphate in the coordination of phospholipid and lipopolysaccharide synthesis in *Escherichia coli*

**DOI:** 10.64898/2026.03.13.711678

**Authors:** Tanner G. DeHart, Elayne M. Fivenson, Vincent de Bakker, Nazgul Sakenova, Thomas G. Bernhardt

## Abstract

The envelope of Gram-negative bacteria like *Escherichia coli* is multilayered with two membranes sandwiching a peptidoglycan cell wall. The inner membrane is a typical phospholipid bilayer whereas the outer membrane is asymmetric with phospholipids in the inner leaflet and lipopolysaccharide (LPS) in the outer leaflet. We recently discovered that inactivation of the conserved peptidoglycan synthesis machinery responsible for cell elongation causes defects in both peptidoglycan and LPS synthesis in *E. coli*. This finding suggests that the isolation of suppressors that rescue the growth phenotype caused by an impaired cell elongation system is an attractive means of identifying factors involved in coordinating the biogenesis of different envelope layers. Here, we report the results of a global, transposon sequencing-based screen for such suppressors. The inactivation of a number of factors including the phospholipid synthesis enzyme PlsX was found to partially suppress the growth defects of a cell elongation mutant. Deletion of *plsX* also conferred increased resistance to CHIR-090, an inhibitor of the committed step of LPS synthesis catalyzed by LpxC, suggesting that loss of PlsX function stimulates LPS synthesis. Evidence is presented that increased CHIR-090 resistance is not mediated by changes in the activity of the proteolytic system (YejM-LapB-FtsH) controlling LpxC turnover. Rather, our results are consistent with a model in which the phospholipid precursor acyl-phosphate produced by PlsX serves as an inhibitor of LpxC to lower the rate of LPS synthesis when phospholipid synthesis capacity is reduced.

**IMPORTANCE:** Over the last several decades, most proteins essential for Gram-negative cell surface assembly have been characterized. However, relatively little is known about how the synthesis of different envelope layers is coordinated to promote uniform surface growth. Here, we report the results of a transposon sequencing-based genetic screen for mutants that suppress defects in the conserved peptidoglycan synthesis machinery responsible for cell elongation. Inactivation of the *plsX* gene encoding a phospholipid synthesis enzyme was found to both suppress the growth defect of a cell elongation mutant and to confer elevated resistance to an inhibitor of lipopolysaccharide synthesis. Our results suggest the attractive possibility that the product of PlsX, acyl-phosphate, may play a regulatory role in coordinating the phospholipid and lipopolysaccharide synthesis pathways.

## INTRODUCTION

Many significant human pathogens such as *Escherichia coli*, *Pseudomonas aeruginosa*, and *Vibrio cholerae* are characterized as Gram-negative based on the staining properties of their cell envelope. Infections with these organisms are becoming increasingly difficult to treat, in part due to the intrinsic antibiotic resistance conferred by the complex organization of their surface (1) . The cell envelope of Gram-negative bacteria is composed of a thin peptidoglycan (PG) cell wall sandwiched between an inner and an outer membrane (2) . The inner membrane (IM) is a symmetrical phospholipid (PL) bilayer that encapsulates the cytoplasm whereas the outer membrane (OM) is asymmetric consisting of an inner leaflet of PLs and an outer leaflet of lipopolysaccharide (LPS) glycolipid (2, 3). The PG layer is built in the periplasmic space between the membranes (4). It along with the OM provides cells with the mechanical stability needed to resist osmotic pressure and prevent cell lysis (2, 5–10) . Over the last few decades most of the essential factors needed for the assembly of the envelope have been identified and characterized. What remains largely unclear is how the biogenesis of the different envelope layers is coordinated to ensure uniform surface growth (11) .

In recent years, defects in the conserved PG synthesis machinery called the Rod system (elongasome) in *E. coli* have been shown to cause a dramatic reduction in LPS synthesis in addition to interfering with proper PG biogenesis (8) . Although the underlying cause of this phenomenon is poorly understood, it indicates that OM assembly and PG biogenesis by the Rod system are linked in some way. The isolation of spontaneous suppressors that rescue the growth defect of cells encoding a hypomorphic allele of a Rod system component [*mreC*(*R292H*)] has therefore proven useful for uncovering new insights into both PG and OM biology. For example, the characterization of such suppressors helped reveal the regulatory mechanisms controlling Rod system activity and identified a role for the OM in cell shape determination (8, 12–14). We therefore reasoned that a global, transposon sequencing-based search for additional suppressors of Rod system defects had the potential to identify new factors controlling envelope assembly, including those responsible for coordinating the synthesis of different envelope layers. Here, we report the results of this screen, which identified a number of genes that suppress the growth defect of the Rod system hypomorph when they are inactivated. We chose to investigate how *plsX* inactivation promotes growth of the *mreC*(*R292H*) mutant because it encodes a PL synthesis enzyme (15), suggesting that further study of the underlying suppression mechanism might reveal a link between PG and/or LPS biogenesis with PL synthesis.

PlsX is involved in a two-step pathway needed to convert long chain fatty acids in the form of acyl-acyl carrier protein (acyl-ACP) conjugates into the essential PL precursor lysophosphatidic acid (LPA) (15) (**Fig. 1**). PlsX converts acyl-ACP into the intermediate acyl-phosphate (acyl-P). The second step is catalyzed by PlsY, which transfers the acyl chain from acyl-P to glycerol-3-phosphate (G3P) to form LPA. PlsC adds a second acyl chain to LPA to form phosphatidic acid (PA), the intermediate to which headgroups are added to produce mature PLs (16). Most bacteria only encode the PlsXY pathway for LPA synthesis (15, 16). However, *E. coli* and many other gamma-proteobacteria encode a second LPA synthesis enzyme called PlsB that is homologous to the PL synthesis enzyme used by eukaryotes (16) (**Fig. S1**). PlsB directly transfers an acyl group from acyl-ACP to G3P to form LPA (**Fig. 1**). In these organisms, PlsB is essential whereas the PlsXY pathway is dispensable (17). Individual deletions of *plsX* or *plsY* are tolerated, but strangely the simultaneous inactivation of both genes is lethal (17). Why *E. coli* and related bacteria have maintained the PlsXY pathway given that they also produce PlsB has remained unclear but may be due to their role in preventing the toxic accumulation of acyl-ACP along with the possible acquisition of other alternative functions for these enzymes (17, 18) .

**Figure 1:**
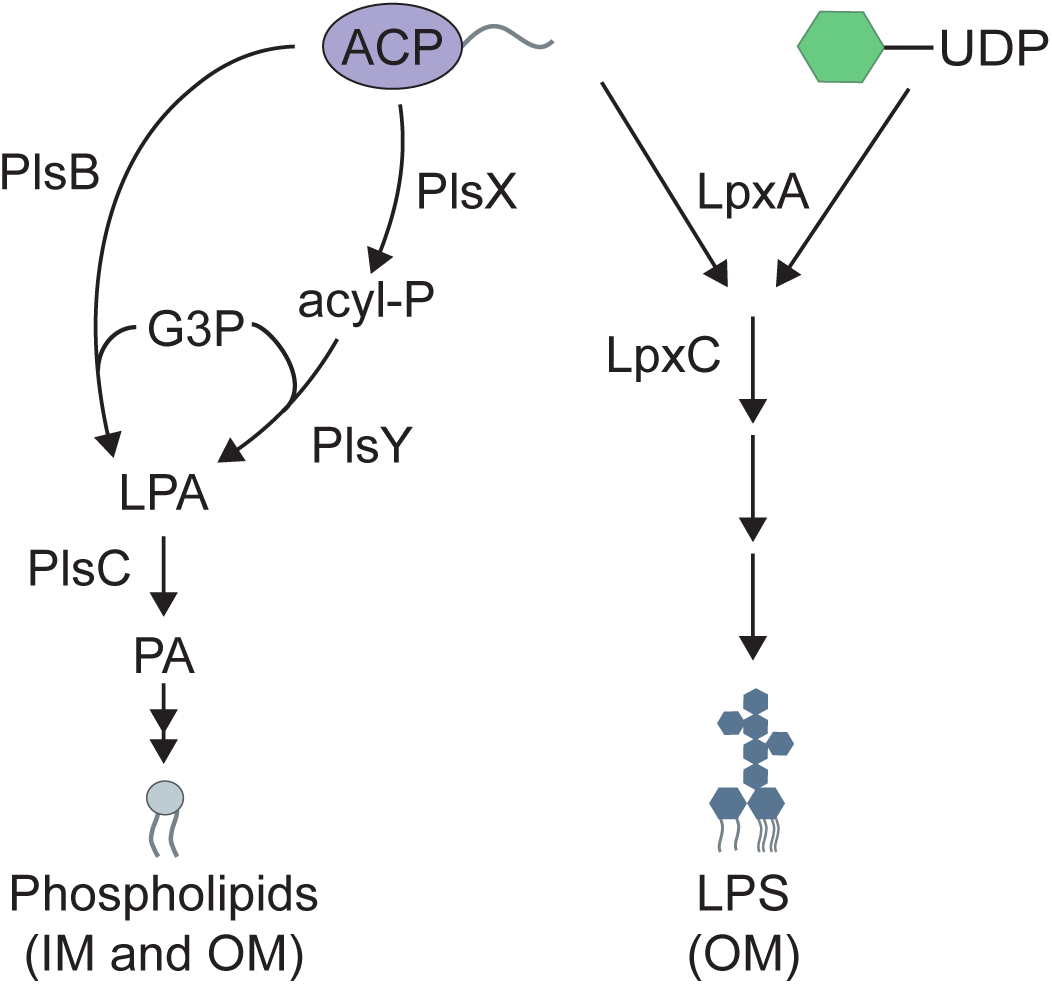
Pathways for PL and LPS biosynthesis. Shown are simplified diagrams depicting the PL and LPS synthesis pathways. Note that long chain acyl-ACPs (C16-18) are used by the PL synthesis pathway while medium chain acyl-ACPs (C12-14) are used by the LPS synthesis pathway. UDP-Glc*N*Ac (green hexagon) is necessary for the synthesis of LPS as well as the PG cell wall. G3P = glycerol-3-P, acyl-P = acyl-phosphate, LPA = lysophosphatidic acid, PA = phosphatidic acid. See text for details.

Deletion of *plsX* but not *plsY* was found to partially suppresses the growth and morphology defects of the Rod system hypomorph. Inactivation of PlsX also conferred increased resistance to CHIR-090, an inhibitor of the committed step of LPS synthesis catalyzed by LpxC (19), suggesting that PlsX activity is somehow antagonistic to LPS synthesis. Evidence is presented that increased CHIR-090 resistance is not mediated by changes in the activity of YejM, a regulator of LpxC proteolysis (20–25) that has also been found to interact with PlsY (20) and thus may link PL and LPS synthesis. Rather, our results are consistent with a model in which acyl-P serves as an inhibitor of LpxC to lower the rate of LPS synthesis when the capacity for phospholipid synthesis is reduced. This proposed regulatory activity of acyl-P is another potential reason why the PlsXY pathway has been maintained in gamma-proteobacteria despite it being non-essential for PL synthesis.

## RESULTS

### A global screen for suppressors of Rod system defects

The Rod system is a PG biogenesis machine that promotes cell elongation and maintains rod shape (4) . It is composed of the RodA-PBP2 PG synthase complex, three membrane proteins of poorly understood function (RodZ, MreC and MreD), and filaments of the actin-like MreB protein that orient PG synthesis by the system. We previously identified hypomorphic variants of the MreC component of the complex in *E. coli* that, like deletion alleles of Rod system genes, caused loss of rod shape and a conditionally lethal growth defect (12, 13). The mutants survive on minimal medium but are not viable on rich medium (lysogeny broth, LB). Selections for spontaneous suppressors of two different *mreC* mutants, *mreC(R292H)* or *mreC(G156D)*, that restored growth under non-permissive conditions and at least partially restored rod shape yielded mutants encoding hyperactive variants of RodA and PBP2 (13). These suppressors provided important insight into Rod system regulation and implicated MreC in the activation of PG synthesis by the complex (13, 14). Another class of suppressors mapped to genes encoding the LpxC proteolytic system (YejM-LapB-FtsH), revealing that Rod system defects cause the aberrant degradation of LpxC and reduced LPS synthesis (8). Characterization of this subset of *mreC* suppressors has proven useful in uncovering the mechanism by which LPS synthesis is regulated (21).

The success of the selections for spontaneous suppressors of *mreC* defects in providing new insights into cell envelope biogenesis motivated us to perform a global, transposon sequencing-based screen for the rapid identification of additional *mreC* suppressors. Because they grow poorly even under permissive conditions, it is difficult to generate transposon mutant libraries of *mreC* mutants with a density of insertions sufficient for effective transposon sequencing (Tn-Seq) analysis. We therefore took advantage of the dominant negative activity of the *mreC(R292H)* allele when it is overexpressed from a plasmid (12). A high-density transposon library was generated in a wild-type background (MG1655 derivative) and then transformed with a plasmid encoding either *mreC(WT)* or *mreC(R292H)* under control of an IPTG-inducible promoter (P*_tac_*). The transformed libraries were then plated on LB supplemented with IPTG to induce expression of the *mreC* allele. Overexpression of *mreC(WT)* is tolerated under these conditions, but induction of *mreC(R292H)* is lethal because the dominant negative allele disrupts Rod system function. Most mutants in the library survive when *mreC(WT)* is produced, whereas only mutants with transposon insertions that suppress the growth defect caused by Rod system inactivation will survive in the population overexpressing *mreC(R292H)*. Thus, insertions that suppress Rod system defects were identified as those that became significantly enriched in the population of cells expressing *mreC(R292H)* versus those producing *mreC(WT)* (**Fig. 2 and Fig. S2**).

**Figure 2:**
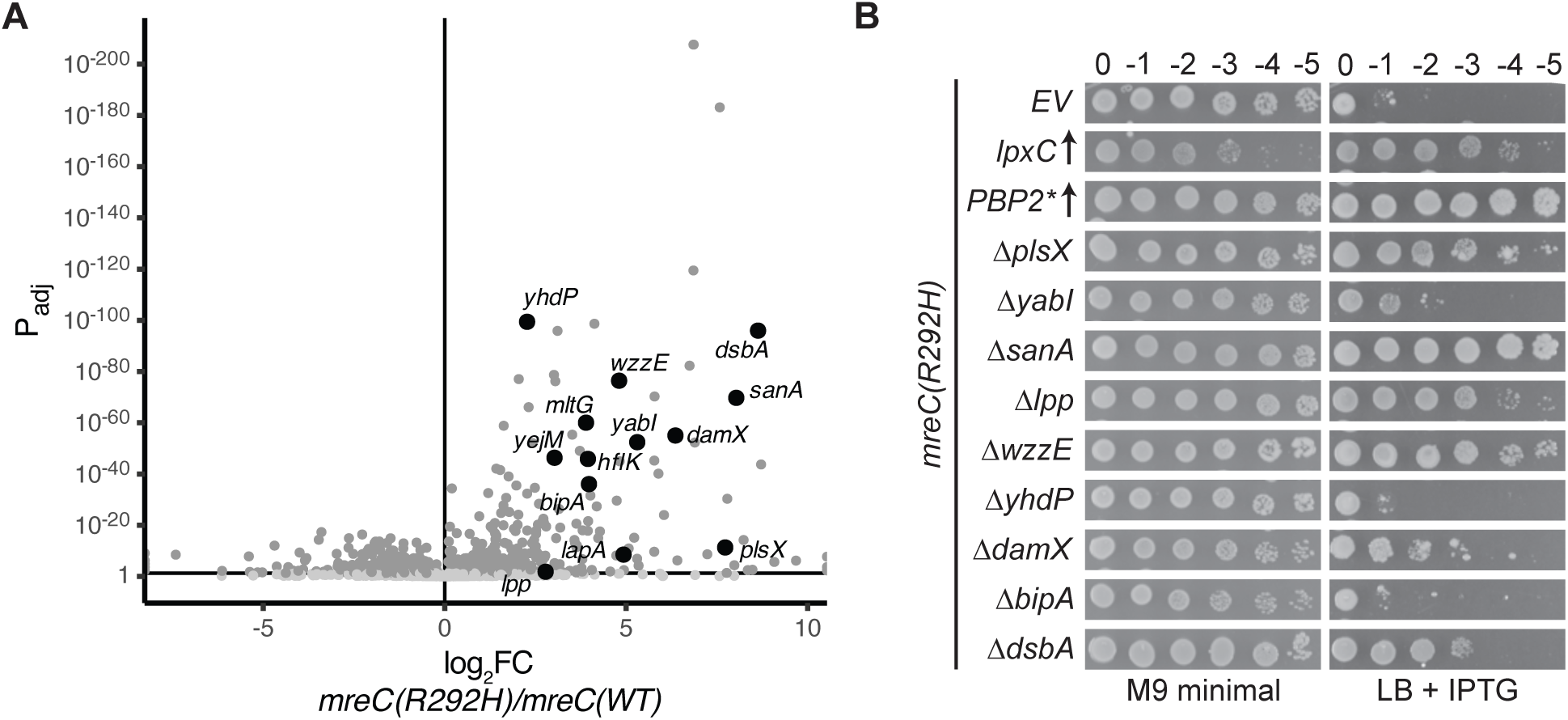
Tn-Seq screen to identify suppressors of the *mreC(R292H)* growth defect. **A.** Volcano plot depicting the fold change (x-axis) and adjusted p-value (y-axis) of transposon insertions into the indicated genes of MG1655 cells expressing *mreC(R292H)* from pPR49 [*P_tac_::mreC(R292H) mreD*] relative to those expressing *mreC(WT)* from pPR11(*P_tac_::mreC(WT) mreD*). Light gray dots indicate a non-significant p-value of > 0.05, while dark gray indicates a significant p-value of < 0.05. Genes bolded in black with labels were among the most significant hits that also have known or predicted roles in cell envelope biogenesis. **B.** Viability of *mreC(R292H)* cells harboring the indicated single deletion mutations. Cells were spotted on LB + 1 mM IPTG at 30°C for 43 hours (right, non-permissive condition) or M9 minimal media + 0.2% casamino acids + 0.2% glucose for 43 hours at 30°C as a control (left, permissive condition). Cells of the *mreC(R292H)* mutant in the first three rows harbor the plasmids pPR66 [empty vector, EV], pPR111 [P*_lac_*::*lpxC*], or pEMF111 [P*_lac_*::*mrdA(L61R) rodA*], respectively.

As an indication that the screen was working as expected, several loci displaying an enrichment of insertions in *mreC(R292H)* expressing cells encoded factors related to those previously implicated in the suppression of Rod system defects. For example, insertions in *yejM*, *lapA* (26), and *hflK* (27) are all expected to impact the activity of the YejM-LapB-FtsH system responsible for LpxC turnover (20–26, 28, 29), and insertions in *mltG* inactivate a PG cleaving enzyme known to antagonize Rod system activity (30) (**Fig. 2 and Fig. S2**). The screening results were further validated for nine additional genes displaying an enrichment of insertions in *mreC(R292H)* overexpressing cells that also encode factors with known or predicted roles in cell envelope biogenesis (*yabI, sanA, wzzE, yhdP, bipA, dsbA, damX, lpp,* and *plsX*) (**Table 1**). To this end, deletions of each gene were individually introduced into a mutant with *mreC(R292H)* at the native locus and the resulting strains were tested for suppression of the growth defects caused by the *mreC* allele alongside controls (**Fig. 2**). Six of the nine deletions restored growth the of the *mreC(R292H)* mutant cells on LB medium, indicating that the Tn-Seq screen was effective at identifying new *mreC(R292H)* suppressors.

### Inactivation of PlsX confers increased A22 and CHIR-090 resistance

Previously isolated suppressors were found to promote growth of *mreC(R292H)* mutants either by activating the Rod system or by increasing LPS synthesis (8, 12, 13). To classify the newly identified suppressors from the Tn-Seq screen, we tested deletion mutants of the genes in question for their ability to confer increased resistance to A22, an antagonist of the Rod system (31), or CHIR-090, an inhibitor of the committed step of LPS synthesis catalyzed by LpxC (19). Most of the deletion mutants that suppressed the growth defect of *mreC(R292H)* cells also conferred increased resistance to A22. In many cases, the increased A22 resistance observed was comparable to that mediated by the overproduction of an activated variant of PBP2 [PBP2(L61R), PBP2*] (13) (**Fig. 3A**), indicating that these deletions are likely to either promote Rod system activity or to ameliorate the detrimental effects caused by the loss of rod shape. To distinguish between these possibilities, the morphology of *mreC(R292H)* cells harboring the most potent of the suppressor mutations was assessed (**Fig. 3B and Fig. S3**). Deletion of *plsX* resulted in a partial restoration of cell shape in the *mreC(R292H)* background (**Fig. 3B-C and Fig. S3**). Little change in cell shape, if any, was observed in *mreC(R292H)* cells harboring deletions in the other genes tested (**Fig. 3B**). Therefore, many of the suppressors isolated are likely to promote the growth of cells with Rod system defects by means other than Rod system activation. Only PlsX inactivation promoted the restoration of rod shape in a manner indicative of Rod system activation.

**Figure 3:**
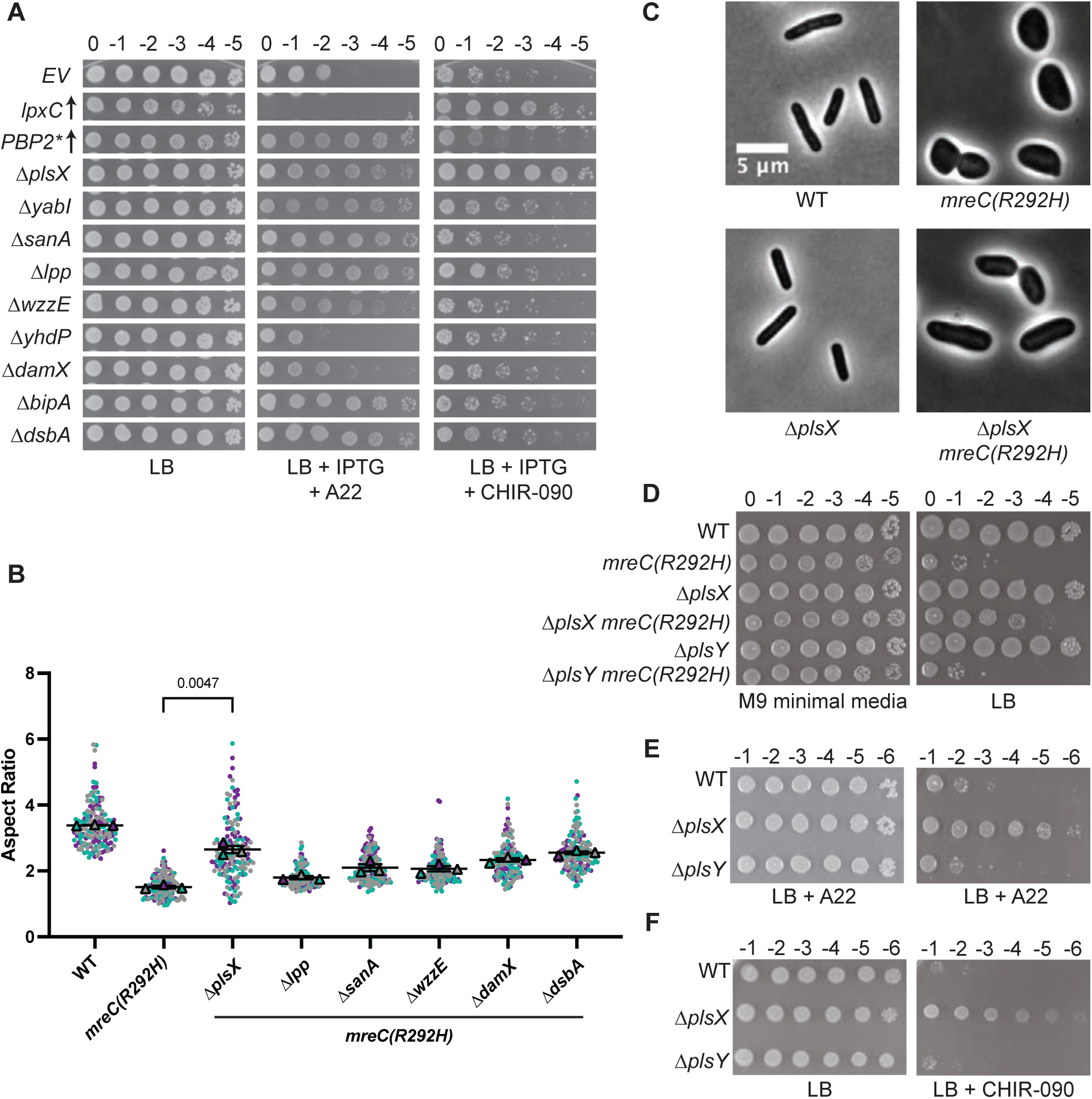
Properties of the *mreC(R292H)* suppressors. **A.** Dilutions of MG1655 (WT) cells with the indicated plasmids or deletions were spotted on LB with or without A22 (2.5 µg/mL) or CHIR-090 (0.05 µg/mL) as indicated. The plates also contained 1 mM IPTG to induce gene expression in the plasmid containing cells. Strains in the first three rows harbor the plasmids pPR66 [empty vector, EV], pPR111 [P*_lac_*::*lpxC*], or pEMF111 [P*_lac_*::*mrdA(L61R) rodA*], respectively. Plates were incubated for 16 hours at 37°C prior to imaging. **B.** Cells of the indicated strains were imaged by phase contrast microscopy and their aspect ratio (length/width) was measured. Replicates for each sample are shown as purple, teal, and gray circles. Each replicate contains approximately 50 cells, for a total of approximately 150 unique cells per strain. Aspect ratio was calculated by determining length and maximum width of individual cells in phase contrast images. The mean of each replicate is shown as a triangle in the respective color of that replicate. An unpaired t-test with Welch’s correction was performed to determine the statistical difference between the means of the *mreC(R292H)* strain and the *ΔplsX mreC(R292H)* strain. The P-value resulting from this analysis is shown above the graph. **C.** Representative micrographs depicting cell shape in MG1655, Δ*plsX*, *mreC(R292H)* and *ΔplsX mreC(R292H)* cells. Cells shown are from the analysis in **B**. **D.** Cells of the indicated strains were spotted on the indicated media and imaged as in Figure 2B. **E-F.** Dilutions of MG1655 (WT) cells with the indicated deletions were spotted on LB with or without A22 (2.3 µg/mL) or CHIR-090 (0.075 µg/mL) as indicated.

Another unique feature of the *plsX* deletion among the suppressors was that it was the only mutant to confer increased resistance to LPS synthesis inhibition by CHIR-090, a phenotype suggestive of LPS synthesis activation (**Fig. 3A**). Accordingly, the level of resistance observed was comparable to that induced by the overexpression of *lpxC* from a plasmid (**Fig. 3A**). Importantly, inactivation of PlsY, the enzyme catalyzing the next step in the PlsXY branch of the PL synthesis pathway (**Fig. 1**), did not suppress the *mreC(R292H)* growth defect or increase resistance to A22 or CHIR-090 (**Fig. 3D-F**). Also, mutants lacking PlsX were not observed to be generally resistant to antibiotic treatments in a prior chemical genetic analysis (32) nor did they display enhanced resistance to other membrane synthesis perturbations such as triclosan treatment (**Fig. S4**). Therefore, the genetic results thus far indicate that deletion of *plsX* but not general defects in the PlsXY pathway promotes Rod system activity and is likely to enhance LPS synthesis.

### Complex effects of the plsX deletion allele

The *plsX* gene is located upstream of the fatty acid biosynthesis genes *fabH*, *fabD*, *fabG*, *acpP*, and *fabF* (33) (**Fig. 4A**). A promoter for the expression of these downstream genes is present at the 3’ end of *plsX* (34). We therefore constructed our *plsX* deletion such that only the first 248 codons of the reading frame were removed, leaving the *fabH* promoter intact (**Fig. 4A**). Nevertheless, to be sure that the phenotypes observed for the deletion were due to the inactivation of PlsX and not an effect on the expression of the downstream genes, we performed complementation tests with *plsX* expressed from a plasmid. Expression of *plsX* in *trans* was only partially able to restore a growth defect to *mreC(R292H) ΔplsX* cells (**Fig. 4B**). Additionally, the increased A22 resistance conferred by the *plsX* deletion was not complemented to normal sensitivity by plasmid-borne *plsX* (**Fig. 4C**). However, the enhanced resistance to CHIR-090 was fully complemented by *plsX* expressed in *trans* (**Fig. 4D**). Thus, the *ΔplsX* allele has complex effects on cell survival when the Rod system is defective. In the case of A22 resistance, the effect appears to principally be due to changes in the expression of downstream genes. However, the partial complementation observed in the context of *mreC(R292H) ΔplsX* cells by plasmid-based *plsX* likely indicates that both the direct effect of the *plsX* deletion on LPS synthesis, as inferred from the increased CHIR-090 resistance, and the indirect effects of the *ΔplsX* allele on downstream gene expression contribute to its ability to suppress the growth defect of *mreC(R292H)* cells. Given the complexity underlying the ability of the *ΔplsX* allele to suppress Rod system defects, we focused the remainder of this study on understanding how PlsX inactivation promotes increased CHIR-090 resistance.

**Figure 4:**
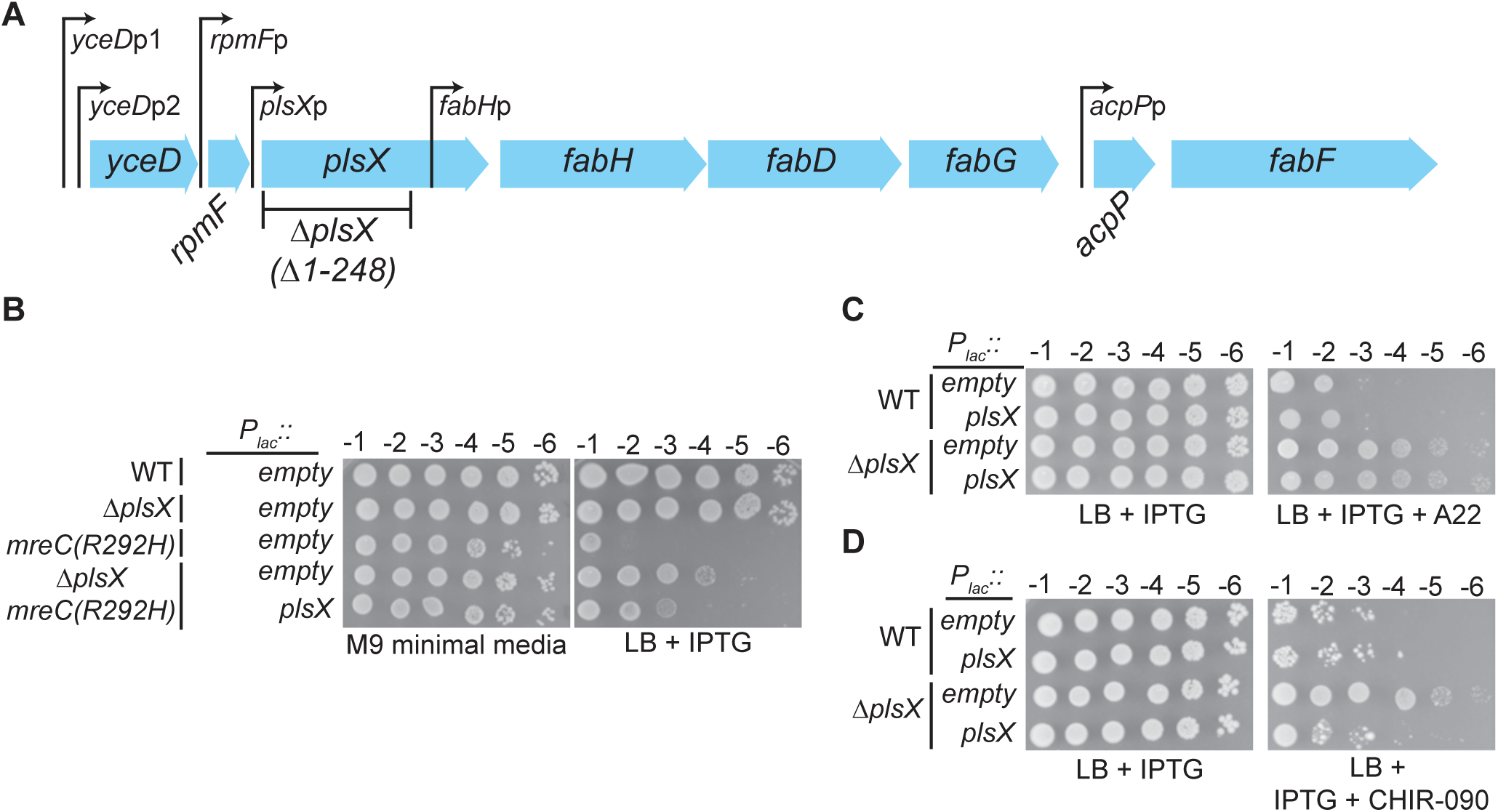
Complex effects of the *plsX(Δ1-248)* deletion allele. **A.** Organization of the Fatty Acid Biosynthesis (FAB) operon. Known promoters are shown as previously described (33). The transcriptional unit of each known promoter is as follows: *yceD*p1 and *yceD*p2: *yceD-rpmF* (53) ; *rmpF*p: *rpmF-plsX-fabHDG* (34, 54); *plsX*p: *plsX-fabHDG* (55); *fabH*p: *fabHDG* (34); *acpP*p: *acpP* (56). **B-D.** Dilutions of the indicted cells harboring pPR66 [empty vector] or pEMF191 [P*_lac_*::*plsX*] were spotted on the indicated media. **B.** Plates were incubated at 30°C for 40 hours (M9) or 21 hours (LB) before imaging. **C.** IPTG (1 mM) and A22 (2.5 µg/mL) were included in the media as indicated. Plates were imaged after growth at 37°C for 16 hours. **D.** IPTG (1 mM) and CHIR-090 (0.075 µg/mL) were included in the media as indicated. Plates were imaged after growth at 37°C for 21 hours.

### The PlsY-YejM interaction is not required for increased CHIR-090 resistance in ΔplsX cells

An attractive model for how the loss of PlsX function might enhance LPS synthesis to confer increased CHIR-090 resistance was suggested by a prior proteomic analysis that detected an interaction between PlsY and YejM (20), the essential regulator of LpxC proteolysis by the FtsH protease and its adapter LapB(YciM) (20–26, 28, 29). PlsX supplies the acyl-P substrate for PlsY. We therefore hypothesized that the absence of acyl-P in *ΔplsX* cells might modulate the PlsY-YejM interaction to affect LPS synthesis. To investigate this possibility, we first confirmed the PlsY-YejM interaction using the POLAR two-hybrid assay (35) (**Fig. 5A**). This cytological assay detects protein-protein interactions in *E. coli* via the ability of a bait protein associated with a polar focus of heterologously expressed *Caulobacter crescentus* PopZ to recruit a prey protein to the cell pole. A GFP-YejM fusion was constructed with an H3H4 domain to target it to polar PopZ foci. Cells expressing this fusion along with a control prey membrane protein fused to mScarlet showed a diffuse mScarlet signal around the periphery, indicating the lack of an interaction between the control prey and polarly localized H3H4-GFP-YejM (**Fig. 5A**). However, when PlsY-mScarlet was used as a prey, it displayed colocalization with H3H4-GFP-YejM indicative of a PlsY-YejM interaction (**Fig. 5A**).

**Figure 5:**
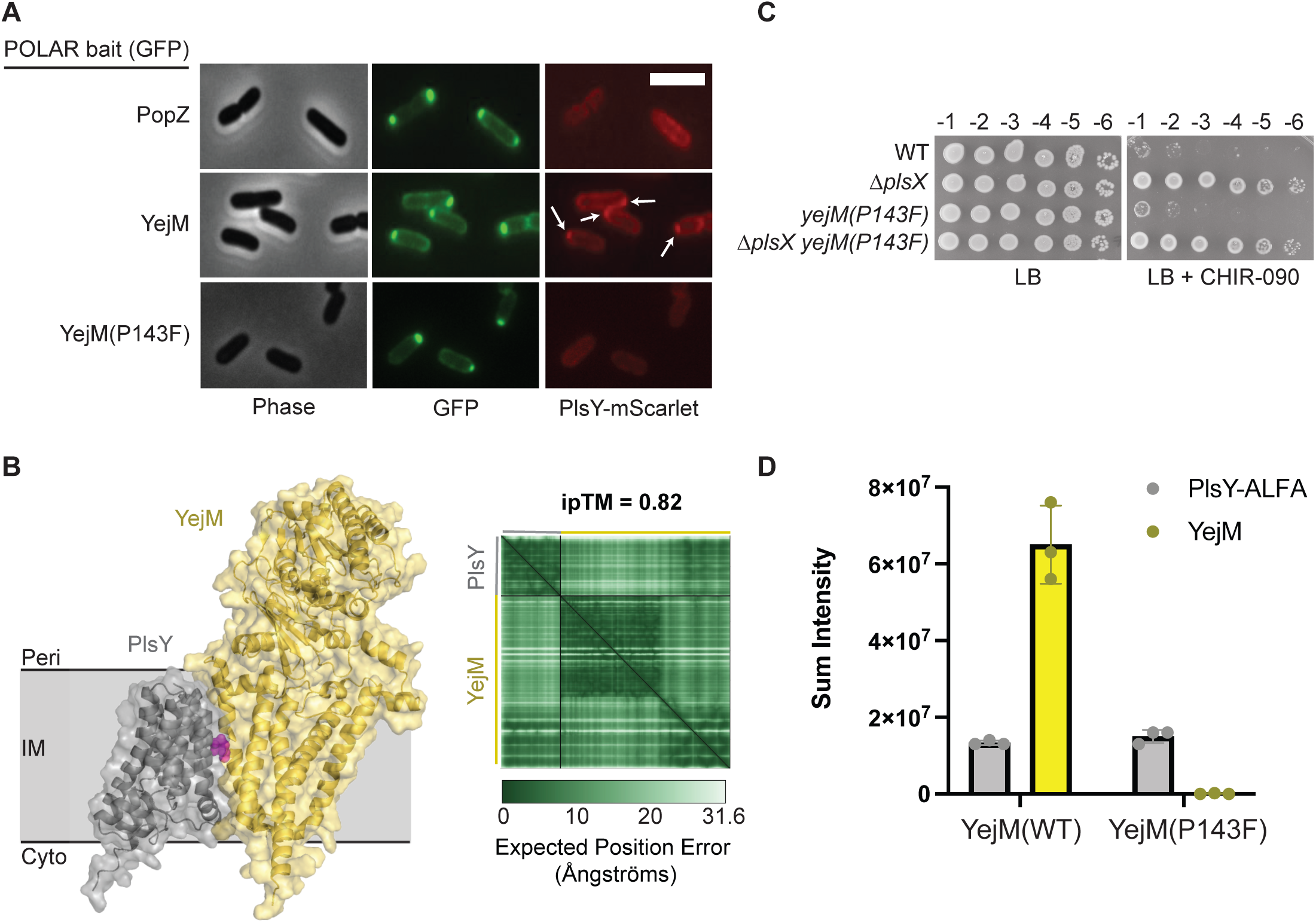
The PlsY-YejM interaction is not required for increased CHIR-090 resistance in Δ*plsX* cells. **A.** POLAR two-hybrid assay demonstrating the interaction of PlsY and YejM can be broken by a single amino acid substitution. Each strain contains a pTGD32(PlsY-mScarlet) “prey” fusion, while also containing either a control “bait” consisting of residues 2-55 of *Pseudomonas aeruginosa* PBP1b fused to H3H4-GFP (pEMF55) (top), or H3H4-GFP-YejM(WT) (pEMF35) (middle), or H3H4-GFP-YejM(P143F) (pTGD73) (bottom). Scale bar = 3 µm. **B.** An AlphaFold 3 model depicting the PlsY-YejM interaction within the inner membrane (IM) oriented with the periplasm (Peri) above and cytoplasm (Cyto) below the IM, respectively. PlsY is shown in gray while YejM is shown in yellow. The P143 residue, mutated to F in the non-interacting variant, is shown in magenta. A Predicted Alignment Error plot depicting the confidence of the PlsY-YejM(WT) interaction is shown to the right. **C.** Cells of the indicated strains were spotted on either LB (left) or LB + 0.075 µg/mL CHIR-090 (right). Plates were incubated for 21 hours at 30°C before imaging. **D.** Sum intensity of PlsY-ALFA and YejM peptides detected in the PlsY-ALFA pulldown elution. Results for biological triplicates are shown.

To test the importance of the PlsY-YejM interaction in modulating CHIR-090 sensitivity in response to PlsX inactivation and for cell growth in general, we sought to identify amino acid substitutions that disrupt the interface. AlphaFold 3 (36) was used to generate a high confidence model of the PlsY-YejM complex, which is predicted to have an interface within the membrane connecting PlsY to the essential transmembrane domain of YejM (**Fig. 5B**). To identify changes that disrupt the interaction detected using the POLAR system, several mutants encoding variants with amino acid substitutions in the H3H4-GFP-YejM or PlsY-mScarlet fusions were generated. Only one substitution variant, H3H4-GFP-YejM(P143F), was found to disrupt PlsY-mScarlet recruitment to the cell pole, suggesting that the P143F change in YejM disrupts the PlsY-YejM interface (**Fig. 5A**).

We next tested the effect of the P143F substitution on YejM function when the variant was produced from its native locus. Cells harboring the *yejM(P143F)* allele were viable and did not display any observable defect in colony formation (**Fig. 5C**). To confirm that the P143F substitution in YejM disrupts its interaction with PlsY, an ALFA-tagged fusion of PlsY (PlsY-ALFA) was produced in wild-type and *yejM(P143F)* cells. Co-purification of YejM was then assessed by mass spectrometry following affinity capture of PlsY-ALFA in each strain (**Fig. 5D**). An equivalent amount of PlsY-ALFA was purified from wild-type and *yejM(P143F)* cells, yet YejM was only detected in the elute from wild-type cells (**Fig. 5D**). These results confirm that the P143F substitution indeed disrupts the YejM-PlsY interaction. Notably, the viability of the *yejM(P143F)* mutant also indicates that the YejM-PlsY interaction interface is not required for the essential function of YejM in controlling LpxC proteolysis (20–25).

To assess whether the YejM-PlsY interaction is required for the increased CHIR-090 resistance displayed by cells lacking PlsX, a *yejM(P143F) ΔplsX* double mutant was generated and its ability to grow on CHIR-090 was compared to control strains (**Fig. 5C**). The CHIR-090 resistance displayed by cells lacking PlsX was unchanged by the presence of the *yejM(P143F)* allele, indicating that the YejM-PlsY interaction is not required for the CHIR-090 resistance phenotype of *ΔplsX* cells. This result was further validated by testing CHIR-090 resistance in *ΔplsX ΔplsY* cells. This mutant combination is normally lethal but has been shown to be tolerated in cells overproducing a cytoplasmic variant of the esterase TesA (^cyto^TesA) (17), which is thought to cleave acyl-ACP molecules to prevent their toxic accumulation in the double mutant. We confirmed this result (**Fig. S5**) and then tested the resulting *ΔplsX ΔplsY* mutant for CHIR-090 resistance. Cells completely lacking PlsY displayed elevated CHIR-090 resistance provided PlsX was also inactivated (**Fig. S5**). Thus, the YejM-PlsY interaction is not required for the CHIR-090 resistance observed upon PlsX inactivation. Importantly, these results also indicate that the YejM-PlsY interaction is not required for the essential function that PlsY performs when PlsX is inactivated. Otherwise, the combination of the *yejM*(P143F) allele with the deletion of *plsX* would have resulted in a lethal phenotype.

### Evidence suggesting that PlsX inactivation enhances LpxC activity

Consistent with the YejM-PlsY interaction being dispensable for the increased CHIR-090 resistance of *ΔplsX* cells, we did not observe a change in LpxC levels in cells lacking PlsX or PlsY as might have been expected if YejM activity were altered by changes in the PlsXY pathway (**Fig. 6**). This result suggested that rather than an increase in LpxC stability it may be LpxC activity that is enhanced when PlsX is inactivated. Accordingly, cells lacking PlsX were hypersensitive to the overproduction of LpxC, suggesting that the toxic accumulation of LPS upon LpxC overproduction occurs at lower levels of the enzyme (**Fig. 6 and Fig. S6**). Curiously, total LPS levels were unchanged in cells lacking PlsX whether or not they also harbored the *mreC(R292H)* allele (**Fig. 6**). Thus, although LpxC activity appears to be increased in *ΔplsX* cells, the overall production of LPS may be limited by additional layers of regulation.

**Figure 6:**
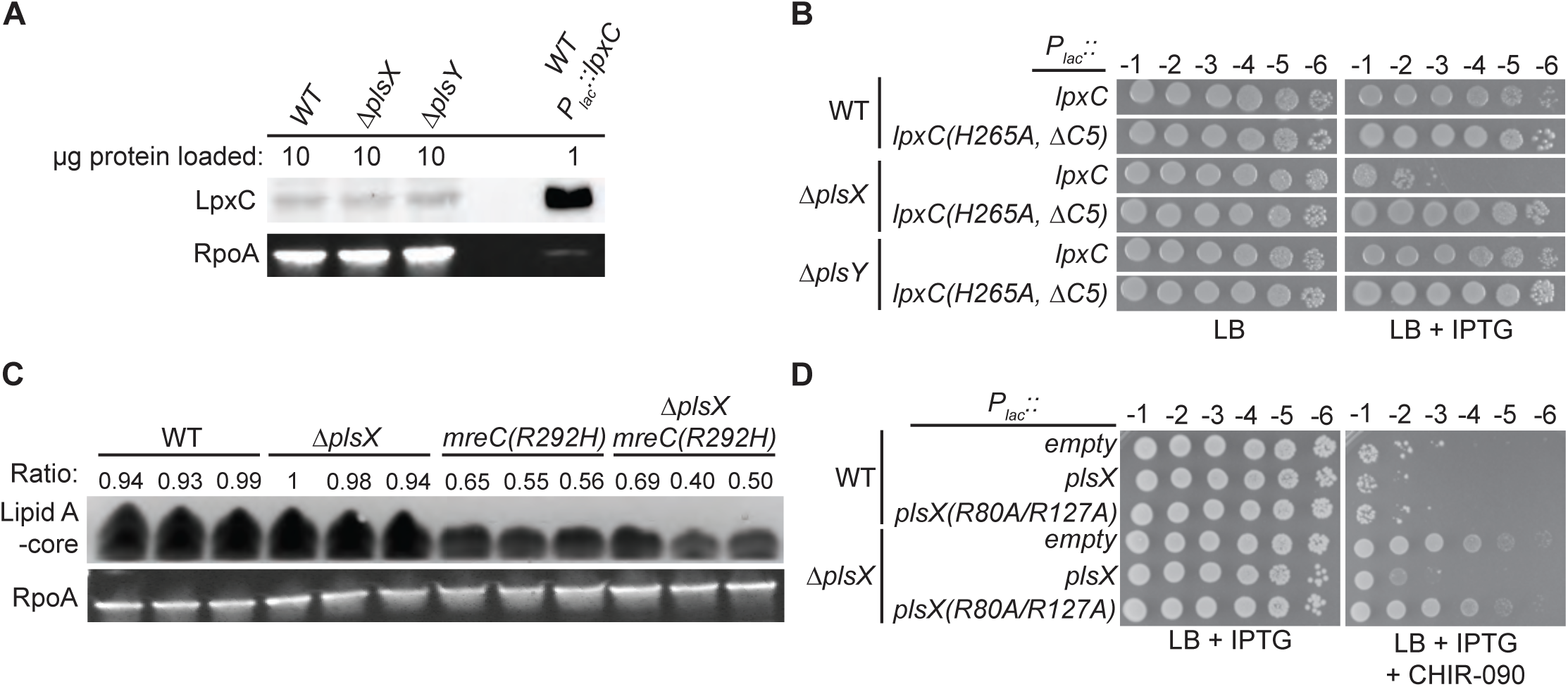
Cells lacking PlsX are hypersensitive to LpxC overproduction. **A.** LpxC was detected by immunoblotting in the indicated strains. RpoA immunoblots were performed to serve as a loading control. **B.** Cells of the indicated strains with plasmids pPR111 [P*_lac_*::*lpxC*] or pPR115 [P*_lac_*::*lpxC(H265A,ΔC5)*] were spotted on LB supplemented with chloramphenicol (25 µg/mL) and IPTG (1 mM) as indicated. The *lpxC(H265A, ΔC5)* allele produces a catalytically inactivated LpxC variant lacking the C-terminal degron. **C.** Silver stain to quantify Lipid A-core levels in MG1655 and *mreC(R292H)* cells with and without *plsX.* The ratio of Lipid A-core normalized by RpoA western blot is shown above each lane. Samples were generated in three separate experiments and then analyzed together with a single silver stain (Lipid A-core) and western blot (RpoA). **D.** Catalytic activity of PlsX is required to complement the CHIR-090 resistance phenotype of Δ*plsX* cells. Cells of the indicated strains and plasmids pPR66 [empty vector], pEMF191 [P*_lac_*::*plsX*], or pTGD160 [P*_lac_*::*plsX(R80A/R127A)*] were spotted on LB with IPTG (1 mM) supplemented with CHIR-090 (0.05 µg/mL) as indicated. Plates were incubated at 37°C for 17 hours before imaging.

How might LpxC activity be modulated by the presence or absence of PlsX? One possibility is that the PlsX protein has a regulatory role independent of its catalytic activity. However, production of a stable, catalytically inactive variant of PlsX failed to restore normal CHIR-090 resistance to *ΔplsX* cells (**Fig. 6**). Thus, these results are consistent with increased CHIR-090 resistance being promoted by the loss of the PlsX product acyl-P not the PlsX protein. Notably, AlphaFold 3 (36) predicts that acyl-P binds within the active site of LpxC in a manner that would be competitive with its substrate (**Fig. S7**). We therefore propose that acyl-P production by PlsX may serve a regulatory role in reducing LPS synthesis at times when PL synthesis capacity is limited (see Discussion).

## DISCUSSION

The Rod system functions to elongate the PG cell wall and maintain rod shape (4). However, inactivation of this system has impacts on envelope assembly that extend beyond problems with PG biogenesis. Defects in the Rod system also cause a dramatic reduction in the synthesis of LPS, which negatively impacts the stress bearing capacity of the OM and its ability to function as a permeability barrier (8). Therefore, selections for suppressors that rescue growth defects caused by Rod system inactivation have been useful for gaining new insights into the regulation of both PG and LPS synthesis (8, 12–14, 21). Here, we used a Tn-Seq based screen to identify a number of new potential suppressors of Rod system defects and validated the suppressing activity of several of them. Because of its role in PL synthesis, we chose to further investigate the mechanism by which PlsX inactivation partially suppresses the growth and shape defects of cells with a poorly active Rod system. Although the *ΔplsX* allele had more complex effects than anticipated with regard to its ability to stimulate Rod system activity and restore rod shape, our results indicate that the loss of PlsX activity has a direct genetic effect on CHIR-090 resistance. Cells with the *ΔplsX* allele are more resistant to the LPS synthesis inhibitor CHIR-090 and this phenotype is complemented by a plasmid-borne copy of *plsX*. This result suggests that similar to other suppressors of Rod system defects, PlsX inactivation stimulates LPS synthesis at the LpxC step targeted by CHIR-090. However, instead of interfering with LpxC proteolysis by the YejM-LapB-FtsH system like previously isolated suppressors (8), inactivation of PlsX appears to promote LpxC activity. To explain this observation, we propose that the acyl-P product of PlsX functions as an LpxC inhibitor such that when PlsX is inactivated, LpxC activity is elevated.

A regulatory role for acyl-P is attractive because it suggests a mechanism by which LPS synthesis can be reduced when problems with PL synthesis arise, especially if G3P were to suddenly become limited (**Fig. 7**). When G3P is abundant, the PlsB pathway is thought to consume most long-chain acyl-ACP molecules in the cell for PL synthesis (15, 17). Under these conditions, PlsX activity and acyl-P production is predicted to be low with the steady state level of acyl-P potentially lowered even further by PlsY activity (**Fig. 7**). However, in G3P limiting conditions, PlsB activity will be reduced, and more acyl-ACP is expected to be consumed by PlsX. Furthermore, the acyl-P generated in this scenario is likely to accumulate given that PlsY activity is also expected to be limited by low G3P levels (**Fig. 7**). Importantly, prior work has shown that the consumption of acyl-ACP by PlsX but not the conversion of acyl-P to LPA by PlsY is essential in cells harboring a PlsB variant with reduced affinity for G3P (17, 37, 38), a condition that would mimic a G3P shortage.

**Figure 7:**
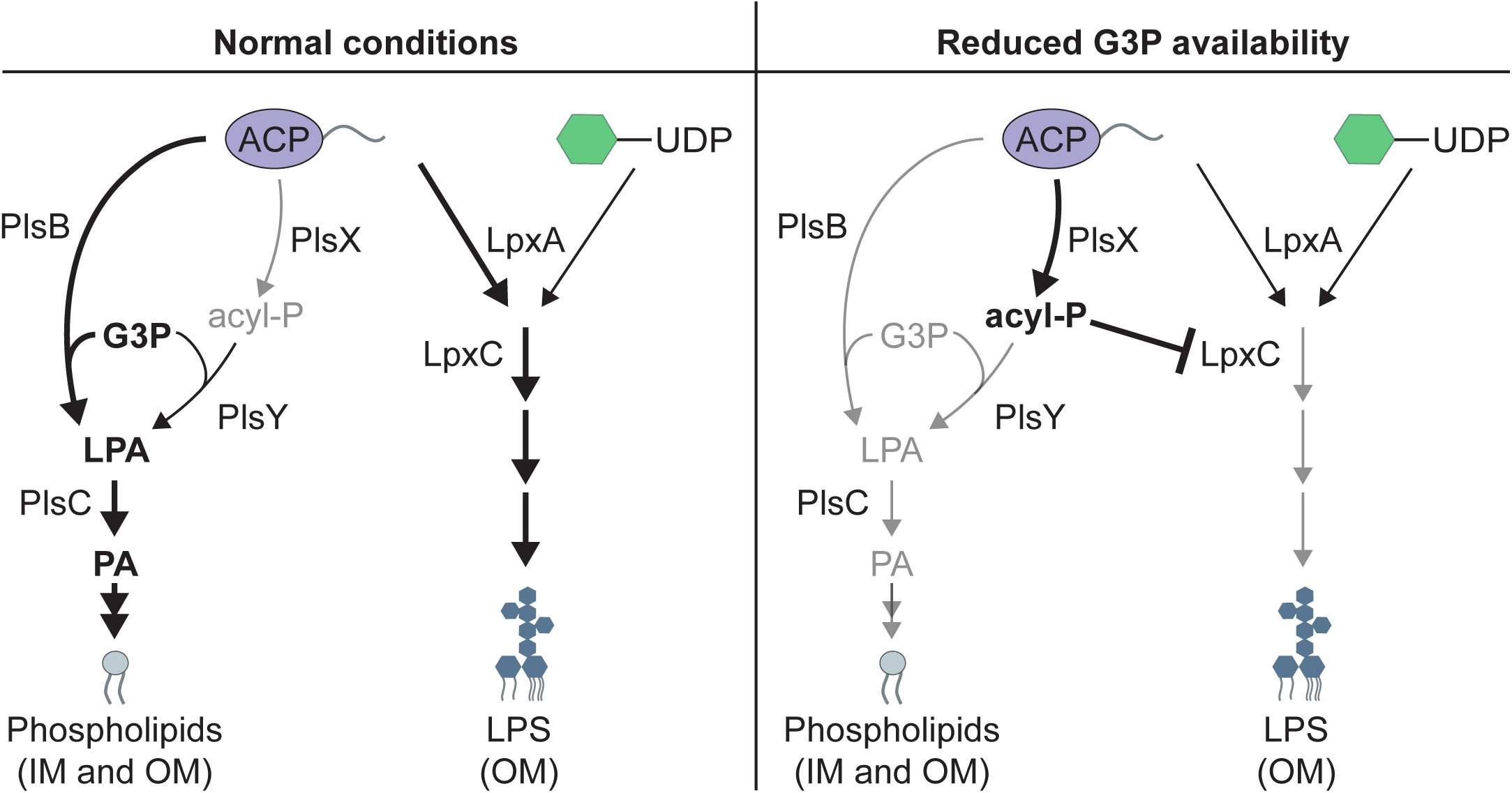
Model for the potential regulatory activity of acyl-P. **Left.** During normal growth G3P levels are expected to be sufficient for PlsB to perform the bulk of PL precursor synthesis. In these conditions, PlsX is expected to compete poorly for acyl-ACP, leading to low steady-state levels of acyl-P, especially if PlsY is actively consuming the acyl-P. **Right.** In conditions were G3P becomes limiting, PlsX is predicted to compete more effectively for acyl-ACP, raising the steady state level of acyl-P. We propose that one consequence of the elevated acyl-P concentration is the inhibition of LpxC to temporarily reduce LPS synthesis and balance it with PL synthesis. See text for details.

The essentiality of PlsX under these conditions has therefore been interpreted to indicate that PlsX activity is not needed for productive PL synthesis when PlsB activity is limited but that it instead functions to prevent the accumulation of toxic levels of long-chain acyl-ACP molecules (17, 18, 37). Indeed, an increase in long-chain acyl-ACPs has recently been observed in cells lacking PlsX (18). Although the consumption of acyl-ACP by PlsX is likely to be important when PlsB is working poorly, we propose that this activity is not its only function. When PL synthesis reduced, it also makes sense for LPS biogenesis to be slowed accordingly, and this reduction could be accomplished by acyl-P inhibiting LpxC as suggested by our genetic results (**Fig. 7**). Thus, acyl-P may serve both as a sink for excess acyl-ACP and a regulatory metabolite that can coordinate PL and LPS synthesis when conditions are in flux.

The acyl-P molecule is labile in aqueous solution, rapidly hydrolyzing to form a free fatty acid and phosphate (39). This instability is an attractive feature for a potential regulatory molecule because it allows for the signal to decay with a short half-life when the adverse conditions stimulating its production are remedied. However, the unstable nature of acyl-P has also made it difficult for us to directly test its proposed role as an LpxC inhibitor. Despite significant effort, we were unable to reliably detect acyl-P in cells whether or not PlsX was overproduced and/or when PlsY was inactivated. Thus, it has not been possible to determine whether acyl-P levels are anticorrelated with LPS synthesis rates as predicted by the model. The instability of acyl-P has also prevented us from reliably measuring the effect of acyl-P on LpxC activity in a purified system. Thus, further testing of the potential regulatory role of acyl-P in coordinating PL and LPS synthesis will require the development of better methods of detecting and working with this unstable molecule.

Given the role of YejM in controlling LpxC proteolysis (20–25) and of PlsY in PL synthesis (15), we became intrigued by the potential role of a YejM-PlsY interaction in coordinating PL and LPS synthesis. We therefore validated the interaction observed in a prior proteomic analysis of YejM interaction partners (20). However, contrary to our expectation, we found that the interaction was not required for the connection between PlsX activity and LPS synthesis that we uncovered nor was it responsible for the essential function of PlsY when PlsX is inactivated. It remains likely that the YejM-PlsY interaction is playing some role in regulating LPS synthesis in response to changes in PL synthesis or vice-versa, but further study is required to define when and how the complex might function in this capacity.

Another enduring mystery surrounds the synthetic lethal phenotype resulting from the simultaneous inactivation of PlsX and PlsY (17). Single deletions of *plsX* or *plsY* will inactivate the PlsXY pathway, yet these mutants are viable. Why then is the double mutant lethal? The ability of ^cyto^TesA overproduction to suppress the growth defect of the *ΔplsX ΔplsY* mutant suggests that acyl-ACP accumulation is at least in part responsible for the synthetic lethal phenotype (17). Accordingly, suppressors that activate PlsB by increasing cellular G3P levels also promote the viability of *ΔplsX ΔplsY* mutants (18). However, loss of PlsX alone has been shown to increase the accumulation of long-chain acyl-ACPs (18). Thus, their accumulation must either increase further to induce lethality when PlsY is inactivated or as recently suggested it is the loss of an as yet unidentified function of PlsY that coupled with the accumulation of long-chain acyl-ACPs induces cell death (18). Identification of this potential unconventional function of PlsY and further exploration of the potential regulatory role of acyl-P in membrane biogenesis will likely be needed to fully understand the lethal phenotype of *ΔplsX ΔplsY* mutants and why the PlsXY pathway has been maintained in *E. coli* and related bacteria that encode PlsB.

Several additional suppressors that rescue the growth defect of mutants defective for Rod system activity were identified in the Tn-Seq screen reported here. Future studies of the suppressing activity of these additional mutants have the potential to shed light on the mechanism(s) connecting Rod system function with OM biogenesis and how the assembly of the different envelope layers is coordinated in Gram-negative bacteria.

## METHODS AND MATERIALS

### Bacterial strain and growth conditions

All strains generated and used in this study are derivatives of MG1655. Strains were cultured in either lysogeny broth (1% tryptone, 0.5% yeast extract, 0.5% NaCl) or modified M9 minimal medium containing 0.5x M9 salts, 0.2% casamino acids, and 0.2% glucose or arabinose, as indicated. Antibiotics were used in the following concentrations unless otherwise noted: chloramphenicol (Cam, 25 µg/mL), kanamycin (Kan, 25 µg/mL), and tetracycline (Tet, 5 µg/mL). All strains, plasmids, and primers used in this study are listed in **Table S1**, **Table S2**, and **Table S3**, respectively. Relevant strain construction information and plasmid construction information can be found in **Table S4** and **Table S5**. All primers were ordered from IDT.

### Strain construction

*Single gene deletions*: Lambda red recombineering was used to create single gene deletions replaced by Kan^R^ as previously described (40–42) . Briefly, the Kan^R^ cassette from pKD13 (or Cam^R^ cassette from pKD3) was PCR amplified with flanking homology to each chromosomal *E. coli* gene slated for deletion. TB10 *E. coli* containing λΔcro-bio were grown overnight at 30°C, back diluted 1:100 into 25 mL of LB broth, and then grown at 30°C to a final OD_600_ of 0.3. Cells were shifted to 42°C for 15 minutes before being placed on ice for 1 hour. Cells were split into two 10 mL aliquots and pelleted at 5000 x g at 4°C for 5 minutes. The supernatant was then removed, and cells were resuspended in 10 mL ice cold glycerol before being pelleted again. Supernatant was removed and cells were resuspended in 1 mL ice cold 10% glycerol and pelleted again. This was repeated for a total of three 1 mL washes in 10% glycerol. Finally, the supernatant was discarded, and cells were resuspended in a final volume of 0.1 mL ice cold 10% glycerol. Electrocompetent TB10 cells prepared above (40 µL) were combined with 2 µL of the PCR product and electroporated at 1.70 kV in a 1 mm electroporation cuvette. Afterwards, 1 mL of LB was swiftly added to the cuvette and the electroporated cells were transferred to a microfuge tube and rolled for 1 hour of outgrowth at 37°C. Finally, these cells were pelleted, resuspended in 200 µL, and plated on LB agar with 25 µg/mL Kan to select for successful replacement of the gene of interest with the Kan^R^ cassette. Unless needed for downstream selection, all antibiotic cassettes were then cured using pCP20 as previously described(41, 43) to generate deletions of chromosomal genes replaced by only a FRT scar.

*Integration of CRIM vectors.* Conditional-Replication, Integration, excision, and Retrieval (CRIM) plasmids were integrated into the chromosome of TB28 cells harboring pTB102 as previously described(44, 45). In brief, an overnight culture of TB28/pTB102 was back diluted 1:100 into fresh LB and grown without antibiotics at 30°C until an OD_600_ of 0.3. The culture was then shifted to 42°C for 15 minutes and afterwards placed on ice for 1 hour. Cells were pelleted at 5,000xg in a tabletop centrifuge before the supernatant was removed and the cells were resuspended in 1/10^th^ the original volume of TSB (LB broth plus 50 µM MgCl_2_, 5% DMSO, and 10% w/v PEG 8000 MW). Now competent, the TB28/pTB102 cells were transformed with the CRIM vector of interest and plated upon the appropriate antibiotic at 37°C overnight. The following day colonies were streak purified on LB Cam at 30°C to ensure loss of the helper plasmid and PCR validated with the primers “CRIM-P2”, “CRIM-P3”, attHK-P1”, and “attHK-P4” to ensure only a single copy of the CRIM vector was integrated into the chromosome. Using these primers, PCR amplification of a single integration of the CRIM vector at attHK022 should yield two bands: 289 and 824 base pairs. Transfer of integrated CRIM vectors between strains was performed by P1 transduction.

*Allelic exchange*. Allelic exchange was adapted from previously described methods (46) to replace *yejM(WT)* with *yejM(P143F)* on the chromosome of EMF191 cells. Briefly, SM10/pTGD89 (donor) and EMF191 (recipient) cultures were grown overnight. From these cultures, 100 µL of donor and 100 µL of recipient were combined in 200 µL of LB in a microfuge tube for a total of 400 µL. For a donor-only and recipient-only controls, 200 µL of donor or recipient cells were added to 200 µL of LB. All cultures were then incubated at 37°C for 6 hours and then plated at various dilutions on LB containing both 25 µg/mL Cam and 25 µg/mL Kan to select for recipient cells (Kan^R^) with an integrated pTGD89 (Cam^R^) vector from the donor strain. Single colonies were picked from the double selection plate and resuspended in 50 µL of 10 mM MgSO_4_. This suspension was serially diluted and 100 µL of each dilution was plated on LB + 6% sucrose at 30°C to counter select against *sacB* encoded on the integrated pTGD89. This condition selects for the excision of the pTGD89 vector. The following day, single colonies were patched on LB and LB + 25 µg/mL Cam to check for the loss of pTGD89. The *yejM* gene in the Cam sensitive colonies was sequenced to determine which colonies contained the *yejM(P143F)* allele. The presence of a downstream Kan^R^ cassette allowed for the *yejM(P143F)* allele to be transduced into the appropriate genetic backgrounds.

### Molecular biology

All restriction enzymes, reagents, and buffers used in plasmid construction were purchased from New England Biolabs (NEB). All polymerase chain reactions (PCR) were performed using Q5 high fidelity polymerase following NEB’s protocol. All restriction digest reactions were performed using NEB restriction enzymes and CutSmart buffer following NEB’s protocol. All ligations were performed using T4 DNA ligase and T4 DNA ligase buffer following NEB’s protocol. Single nucleotide substitutions and small insertions/deletions were introduced using Kinase-Ligase-Dpn1 (KLD) enzyme mix and buffer from NEB, following manufacturer’s protocol. Plasmids were miniprepped using the QiaPrep Spin Miniprep Kit from Qiagen, following manufacturer’s protocol. All plasmids were sequenced by Plasmidsaurus.

### Transposon sequencing

*Transposon mutagenesis:* Transposon mutagenesis of EMF193 was performed as previously described (47). EMF193 cells were transposon mutagenized using the donor strain MFDλpir/ pSC189 (48). This donor strain requires 300 µM diaminopimelic acid (DAP) supplemention for growth. It delivers the Kan^R^ mariner transposon to the recipient strain via conjugation. Mutagenized cells were selected on LB Kan agar without DAP. The transposon mutagenized libraries were scraped from the LB agar plates, resuspended in LB Kan at an OD_600_ of 10. Glycerol was added at a final concentration of 16% glycerol at aliquots of the library were stored at -80°C. The library (1 mL) was thawed and added to a flask containing 100 mL of LB to make electrocompetent cells which were ultimately transformed with pPR11 [P*_tac_*::*mreC(WT)*] or pPR49[P*_tac_*::*mreC(R292H)*]. Transformant libraries were then scraped, resuspended at an OD_600_ of 10, and stored as for the original library.

*Transposon insertion sequencing.* Transposon insertion sequencing was performed as previously described (47). Genomic DNA was extracted from the final, glycerol stocked, transposon mutagenesis libraries described in the previous section. The genomic DNA was then fragmented via sonication, purified, and modified to add poly-C tails. Transposon-chromosome junctions were amplified via PCR using the following primers:

*PolyG-1st-1*:

*5ʹ-GTGACTGGAGTTCAGACGTGTGCTCTTCCGATCTGGGGGGGGGGGGGGGG-3’*

*pSC189-1st-3*:

*5ʹ-GGCTGACCGCTTCCTCGTGCTTTAC-3ʹ*

The resulting product was then barcoded through a second PCR reaction using:

*pSC189-2nd-1*:

*5ʹ- AATGATACGGCGACCACCGAGATCTACACTCTTTCGGGGACTTATCAGCCAACCTG-3ʹ*

and primers containing a unique barcode for each sample from the Illumina TruSeq Small RNA suite of barcoding primers. The resulting PCR products were quantified via Qubit (Thermo Fischer Scientific), pooled, and separated via gel electrophoresis in a 2% agarose gel. Bands between 200-500 bp were extracted and gel-purified. Pooled, barcoded, libraries were sequenced using the NextSeq 1000/2000 P1 reagent (100 cycles) alongside the custom sequencing primer:

*BTK30_IL_seq_v2*:

*5ʹ-CTTTCGGGGACTTATCAGCCAACCTGT-TA-3ʹ*

The pooled sequencing data was demultiplexed, and adaptor sequences were trimmed, before being mapped back to the *E. coli* MG1655 reference genome. Transposon insertion profiles were visualized using Artemis Genome Browser (49). Transposon sequencing data are available at http://www.ncbi.nlm.nih.gov/bioproject/1437091.

### Viability assays

Overnight cultures of strains of interest were normalized to an OD_600_ of 1.0. After normalization, cultures were serially diluted 1:10 in fresh LB medium. Using a multichannel pipette, 3 µL of a range of each serially diluted culture was plated on agar plates. For Figures 3A and 3D, 5µL was spotted. The composition of each agar plate and range of serial dilution spotted are indicated on each figure. For assays with strains containing expression vectors, in each case the overnight cultures were back-diluted 1:50 into fresh media + inducer and grown for three hours prior to normalization and spotting as indicated in the corresponding figure.

### Quantification of protein levels, immunoblotting, and LPS staining

*Preparation of samples for western blots and silver stains:* Overnight cultures of strains of interest were back diluted to an OD_600_ of 0.025 in 5 mL of LB and grown to an OD_600_ of 0.4. Afterwards, cultures were back diluted to an OD_600_ of 0.025 in 50 mL of LB + inducer (if necessary). Cultures were grown to an OD_600_ of 0.5, or for a set amount of time, as described. The final OD_600_ was measured, and 20 mL of culture was pelleted at 5000 x g for 10 minutes at 4°C before being resuspended at a final OD_600_ of 20 in 1:1 water and 2x Laemmli sample buffer containing 5% β-mercaptoethanol (from BioRad Cat. #1610737; final concentration 1x sample buffer, 2.5% β-mercaptoethanol). Samples for silver stains were instead resuspended in 1x LDS sample buffer (Invitrogen NP0008) containing 4% β-mercaptoethanol. Samples were then boiled for 10 minutes and either used immediately or stored at -80°C, as necessary. Total protein levels were quantified for each sample by using the NI (Non-Interfering Protein) Assay Kit from G-Biosciences, following the manufacturer’s protocol.

*Immunoblots.* Samples (20 µg protein) were run on a 4-20% Mini-PROTEAN TGX Precast Gel (Bio-Rad Cat. #456-1094) at 100V until completion and transferred to a polyvinylidene difluoride (PVDF) membrane. The membrane was then blocked in 1x phosphate-buffered saline containing 0.1% Tween (PBS-T) + 5% milk for 1.5 hours, rocking at room temperature. Afterwards, the membrane was incubated with in PBS-T + 1% milk containing the primary antibody of interest. For anti-LpxC blots, the primary was rabbit anti-LpxC antibody (a gift from the Doerrler lab) at a 1:10,000 dilution for 16 hours, rocking at 4°C. For anti-ALFA blots, the primary was anti-ALFA conjugated to horseradish peroxidase (HRP; Cedarlane Labs Cat. # N1505-HRP(SY)) at a dilution of 1:2000. For RpoA controls, the primary was mouse anti-RpoA (Biolegend Cat. #663104) at a dilution of 1:10,000. The membranes were then rinsed with PBS-T and washed three additional times with PBS-T for 10 minutes. The anti-ALFA-HRP blots were then ready for imaging as described below. The anti-LpxC blots were instead incubated with an anti-rabbit IgG-HRP secondary (Rockland no. 18–8816-33) at a dilution of 1:2,000 in PBS-T + 0.2% milk for two hours. The anti-RpoA controls were incubated with anti-mouse HRP at a dilution of 1:3,000 in PBS-T + 0.2% milk for two hours. Post-secondary incubation, membranes were rinsed with PBS-T and then subjected to 4 washes with PBS-T for 10 minutes each. Following the final wash, all membranes were developed using SuperSignal West Pico Plus chemiluminescent substrate (Thermo Fisher Scientific, Cat. #34577) and imaged using the chemiluminescent channel on the Bio-Rad ChemiDoc MP Imaging System.

*Silver staining:* From the samples prepared above, 50 µL was incubated with 1.25 µL of Proteinase K (NEB P8107S) for 1 hour at 55°C. The proteinase K-incubated samples were then heated at 95°C for 10 minutes. The equivalent of 20 µg total protein for each sample was then run on a 4-12% Criterion XT Bis-Tris Precast Gel (Bio-Rad, Cat. #3540124) at 100 V for 2 hours. LPS was detected using the silver stain method (50) . The gel was fixed overnight at room temperature in a solution of 40% ethanol, 5% acetic acid. The following day, periodic acid was added to a final concentration of 0.7% and incubated for 5 minutes. The fixative plus periodic acid solution was then discarded and the gel washed with water. The gel was washed twice for 30 minutes, followed by a 1-hour wash, for a total of 2 hours, rocking at room temperature. The gel was then incubated in 150 mL of staining solution (0.018 N NaOH, 0.4% NH_4_OH, and 0.667% Silver Nitrate) for 10 minutes, rocking at room temperature. Afterwards, the gel was washed 3x with 200 mL of milliQ water, rocking for 15 minutes at room temperature each time. The washes were removed, and the gel was developed via incubation with developer solution (0.26 mM Citric Acid pH 3.0, 0.014% formaldehyde). When bands were visible, the reaction was quenched by removing the developer solution and adding 100 mL of 0.5% acetic acid. The gel was imaged on the Bio-Rad ChemiDoc MP Imaging System.

### ALFA-pulldown, proteomics, and analysis

*Preparation of cultures for pulldown:* Cultures (1 L) of each strain of interest were grown to an OD_600_ of 0.6 in the presence of 1 mM IPTG to induce the ALFA-tagged protein to be pulled down. Cultures were pelleted at 4200 x g, 4°C, for 15 minutes before being resuspended in 50 mL 1x phosphate-buffered saline (PBS). The 50 mL suspension was then pelleted, supernatant was removed, and the pellet was stored at -80°C.

*Lysis and homogenization:* Pellets from above were thawed on ice and resuspended in 30 mL of Lysis Buffer [1x PBS + 0.001% v/v benzonase (Sigma-Aldrich, Cat. #E1014-25KU), 2 “complete” protease inhibitor tablets (Roche, Cat. #11836170001), and 150 mg egg white lysozyme (GoldBio, Cat. #L-040-25)]. Cells were lysed for 30 minutes, rotating, at 4°C. Lysed cell solution was then passed through a cell disruptor seven times and cellular debris was pelleted at 10,000 x g for 10 minutes in an Eppendorf 5810 R tabletop centrifuge. Supernatant was transferred to ultracentrifuge tubes and ultracentrifuged at 35,000 rpm for 45 minutes at 4°C in a Type 45 Ti fixed-angle titanium rotor (Beckman Coulter). Supernatant was removed, and membrane pellets were homogenized into 3 mL membrane buffer [1x PBS + 1% n-Dodecyl-β-D-Maltopyranoside (DDM) (from Anatrace, Cat. #D310)] using dounce homogenizers (Wheaton, Cat. #357544). A new homogenizer was used for each sample. Afterwards, the homogenized solution was transferred to a 15 mL conical tube and rotated end-over-end for 1 hour at 4°C. Finally, an additional 3 mL of membrane buffer was added to each sample.

*Purification:* ALFA magnetic resin (20 µL, Cedarlane Labs, Cat. #N1515(SY)) was aliquoted into an empty 1.5 mL conical tube — one tube with resin for each sample. Resin tubes were placed on a magnetic rack and the supernatant was removed. The resin was washed a total of two times with 1 mL membrane buffer before being resuspended in 20 µL membrane buffer. The resuspended, washed, resin was then added to the 6 mL homogenized membrane solution, from the above section, and incubated at 4°C tumbling end-over-end for 1 hour. Resin was collected in 1.5 mL tubes on magnetic racks — 1 mL of each sample was added, incubated for 5 minutes, and then the supernatant was removed before another 1 mL was added from the same sample to the same tube. Each sample was collected in an individual tube, and this process was repeated 6 times to pool the resin from the entire 6 mL homogenized membrane solution into a single tube. Afterwards, the resin from each 6 mL homogenized membrane solution should be consolidated into a single, unique, microfuge tube for each sample. Pooled resins were washed 4 times total with wash buffer 1-4, in that order. Wash buffers 1-4 contained 1x PBS + 133 mM NaCl + 0.5%, 0.25%, 0.125%, 0.0625% DDM, respectively. For each wash, 200 µL of the respective wash buffer was added to each resin and rotated end-over-end at 4°C for 5 minutes. Afterwards, microfuge tube containing resin + buffer was placed on a magnetic rack for 2 minutes before the supernatant was removed and the next wash buffer was added to repeat the process. After the final wash, the resin was resuspended in 100 µL of elution buffer (1x PBS + 0.05% DDM, 133 mM NaCl, 200 µM ALFA-peptide) [ALFA-peptide: Cedarlane Labs Cat. #N1515(SY)] and rotated end-over-end for 15 minutes at room temperature. Resins were once again collected on a magnetic rack and the supernatant containing ALFA-tagged proteins and any binding partners was collected and stored at -80°C.

*Gels & Proteomic Analysis:* Total eluted protein (20 µL per sample) was run on a 4-20% SDS-PAGE gel (Bio-Rad, Cat. #456-1096) for 30 minutes at 180 V. The gel was washed 3 times with water, for 5 minutes each before being incubated in Imperial Stain (Thermo Scientific, Cat. #24615) for 2 hours at room temperature. The imperial stained gel was quickly rinsed in water 3 times before an overnight destain in water. The following day protein content in each Alfa-tagged elution sample was imaged using the Spyro Ruby channel on the Bio-Rad ChemiDoc MP Imaging System.

The remainder of the total ALFA elution was sent to Taplin Biological Mass Spectrometry Facility at Harvard Medical School for proteomic analysis. The abundance of PlsY-ALFA and YejM in each sample are reported as sum intensity due to the small size of PlsY and few trypsin cleavages sites yielding disproportionately fewer unique peptides compared to other proteins.

*Analysis of fold-change:* Downstream proteomic analyses were performed in R (v4.4.1) with the package DEP (v1.28.0) using sum intensity values as input. Default settings were used, and the imputation step was performed with the MinProb function implemented in DEP, as missing values were biased towards lower intensities and thus not random. Significance of differential enrichment was tested at thresholds of 0.05 for alpha and 0 for absolute log2 fold changes, and p-values were adjusted by false discovery rate (FDR).

### Microscopy

*Phase contrast:* For phase contrast imaging, cells were fixed using a solution of 2.5% formaldehyde + 0.06% glutaraldehyde for 1 hour at room temperature. Fixed samples were immobilized on 2% agarose pads on 1 mm glass slides, using number 1.5 coverslips. Each coverslip was sealed to the slide using a VALAP mixture of 1:1:1 Vaseline:lanolin:paraffin. Imaging was performed using a Nikon TE2000 inverted microscope using Nikon Elements Acquisition software AR 3.2. Cropping and adjustments were made using Fiji (51) software. Cell length and width, used to calculate aspect ratio, were determined using MicrobeJ (52) in Fiji with a rolling-ball transformation of 35 pixels to subtract background noise.

*Two-hybrid localization:* The POLAR assay was utilized to assess the PlsY-YejM interaction as previously described (35). Individual colonies of the appropriate strain were inoculated into 3 mL of LB + 0.2% glucose, 25 µg/mL Cam, and 5 µg/mL Tet and incubated at 37°C for 2 hours, rolling. Each culture was pelleted, supernatant was discarded, and cells were resuspended in 2 mL of M9 minimal media + 0.2% arabinose, 0.2% casamino acids, and 0.1 mM IPTG and incubated at 37°C for 2 hours, rolling. After this incubation, cells were immobilized on agarose pads as described above. Images were taken on a Nikon Ti inverted microscope with a Plan APO lambda 100×/1.45 oil Ph3 DM lens objective, Lumencore SpectraX LED illumination, Chroma ET filter cubes for GFP (49002) and mCherry (49008), an Andor Zyla 4.2 Plus sCMOS camera, and Nikon Elements 4.30 acquisition software. The microscope slide was kept at 30°C using an environmental control chamber.

## ACKNOWLEDGEMENTS

We would like to acknowledge all the members of the Bernhardt and Rudner labs for their advice and thoughtful comments throughout the course of this work. We also thank Paula Montero Llopis and the other members of the MicRoN (Microscopy Resources on the North Quad) team and Ross Tamaino and members of the Taplin Mass Spectrometry Facility at Harvard Medical School for their expertise, support, consultation, and services. We thank Bill Doerrler for the generous gift of the anti-LpxC antibody. This work was supported by the National Institutes of Health (R01 AI083365 to T.G.B.) and Investigator funds from the Howard Hughes Medical Institute (T.G.B.). E.M.F. was supported in part by a National Science Foundation (NSF) Graduate Research Fellowship award. V.d.B. is supported in part by a Swiss National Science Foundation (SNSF) Postdoc Mobility fellowship (P500PB_225439).

## SUPPLEMENTAL MATERIAL

**Figure S1:**
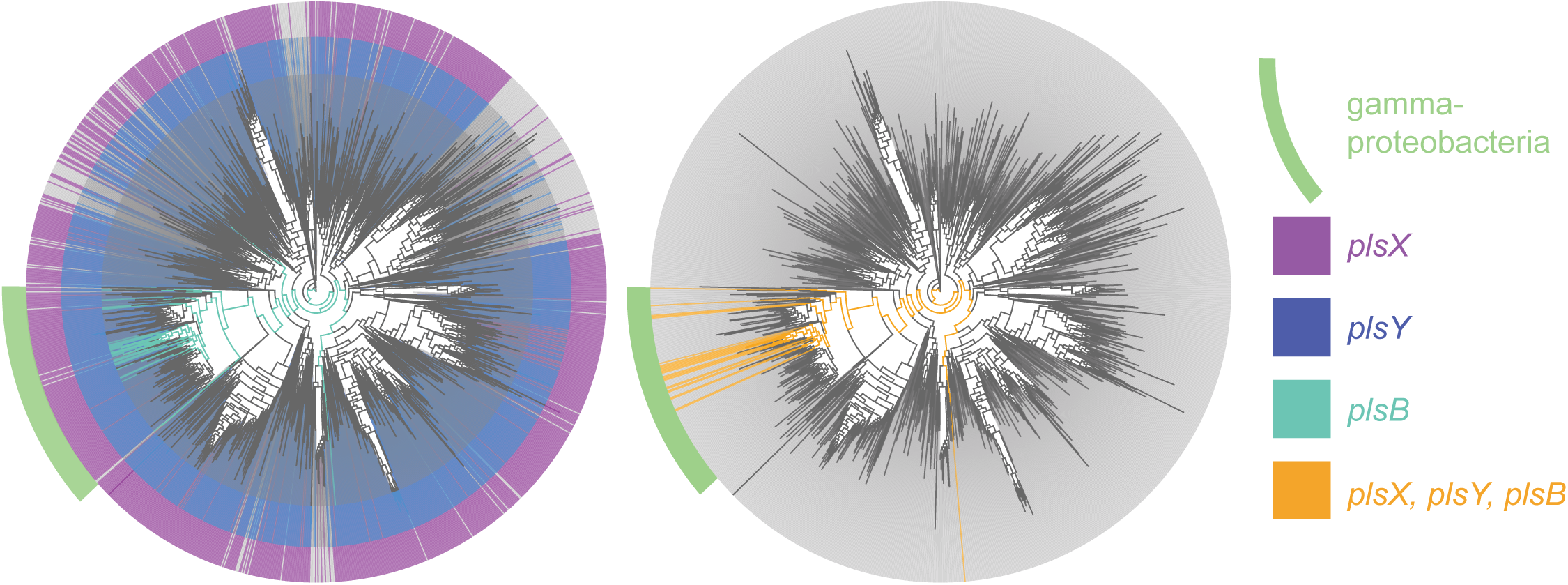
Conservation of phospholipid synthesis genes across bacteria. Each line represents an order of bacteria. Orders within the bacterial class gamma-proteobacteria are shown by the green arc. **Left.** The presence of *plsX*, *plsY*, *plsB* in each genome is represented by purple, blue, or teal, respectively. **Right.** Species that possess *plsX*, *plsY*, and *plsB* are shown highlighted in orange. This figure was made with AnnoTree (1).

**Figure S2:**
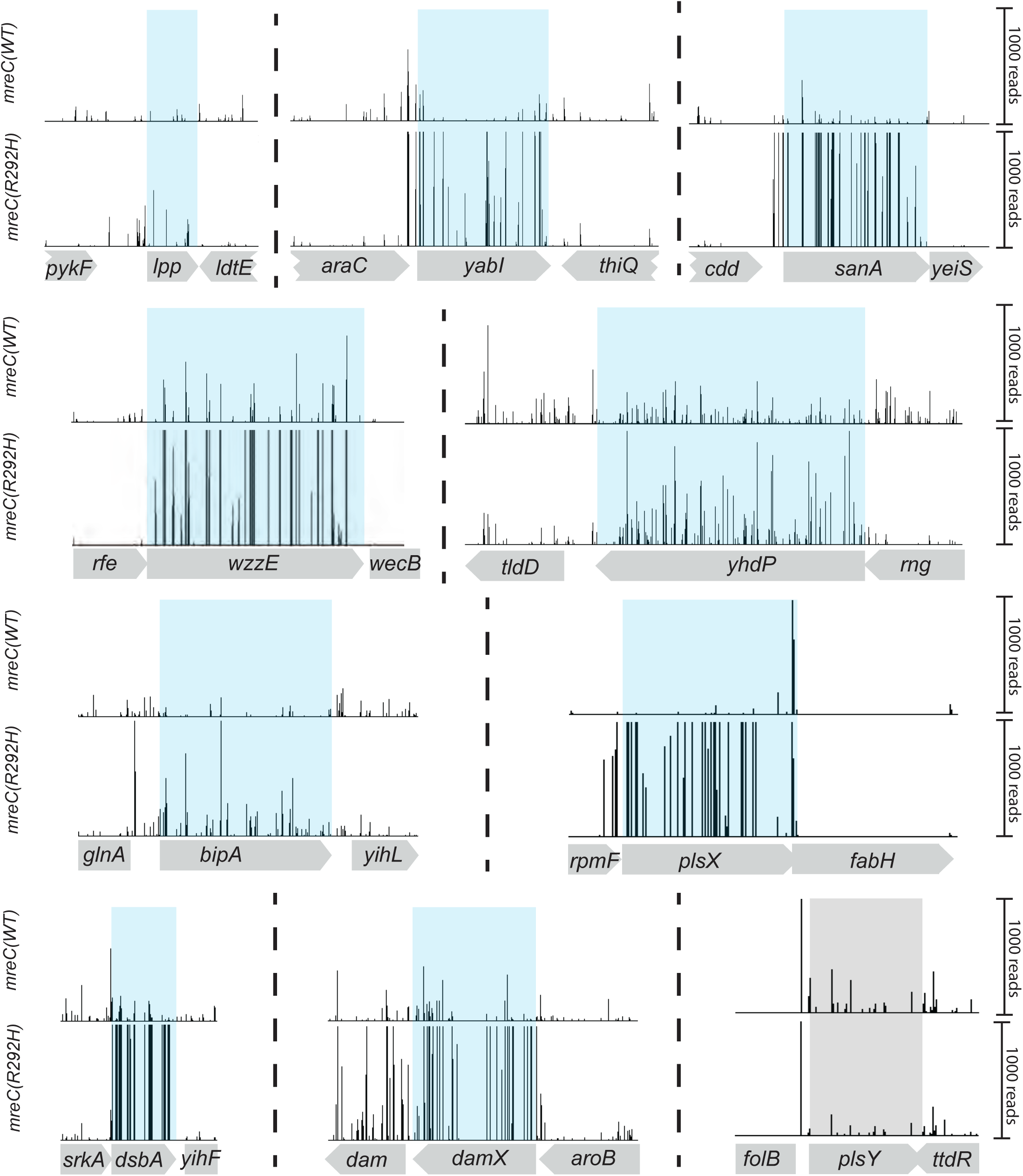
Transposon insertion profiles of genes identified as hits. Tn-Seq insertion profiles for the indicated genes in MG1655 cells expressing *mreC(R292H)* from pPR49 [*P_tac_::mreC(R292H) mreD*] relative to those expressing *mreC(WT)* from pPR11[*P_tac_::mreC(WT) mreD*]. The relative height of each vertical line indicates the frequency of insertions at that site in the population. The profile of *plsY* is highlighted in gray to indicate that it was not called at a hit. It is shown for comparison with the *plsX* profile.

**Figure S3:**
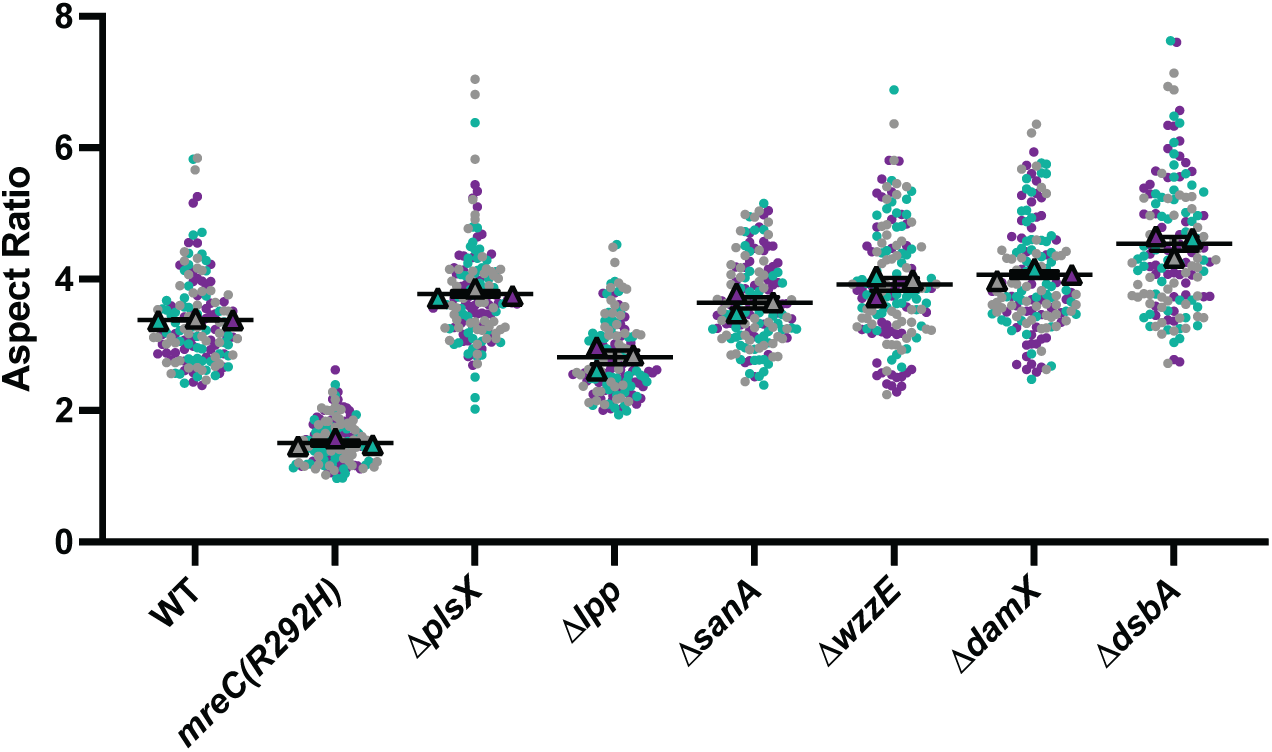
Cell aspect ratio measurements. Cells of the indicated strains were imaged by phase contrast microscopy and their aspect ratio (length/width) was measured as in Figure 3.

**Figure S4:**
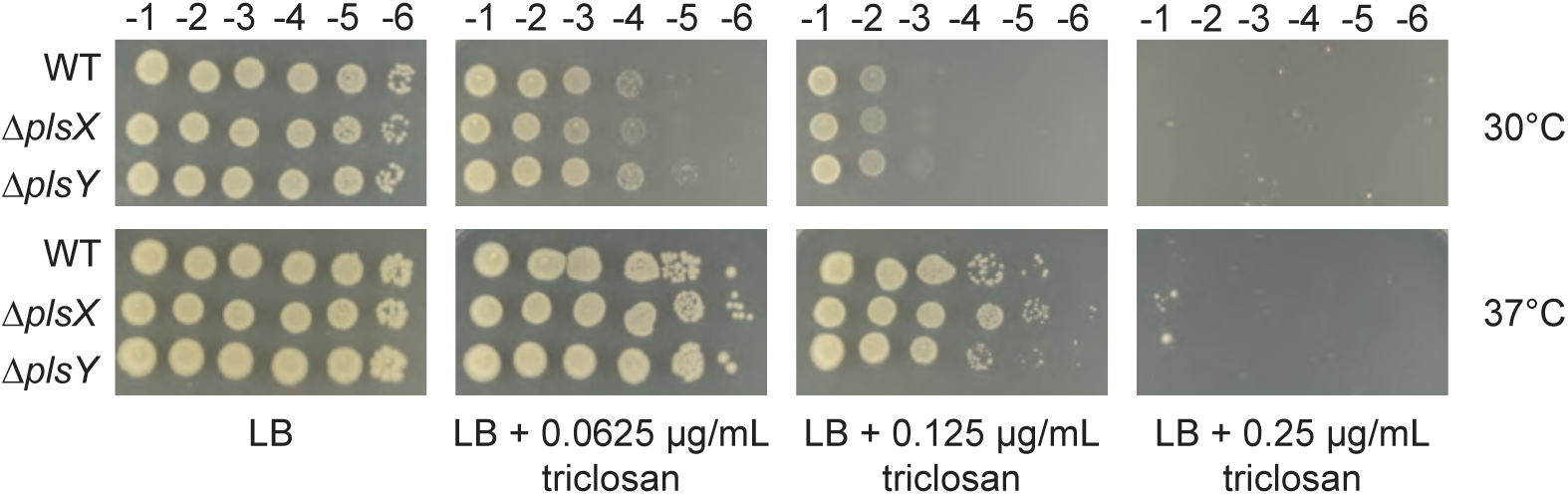
Loss of *plsX* does not confer a resistance to triclosan. Serial dilutions of MG1655, *ΔplsX*, and *ΔplsY* cells were spotted on LB supplemented with various concentrations of triclosan at either 30°C (**top**) or 37°C (**bottom**) for 16 hours.

**Figure S5:**
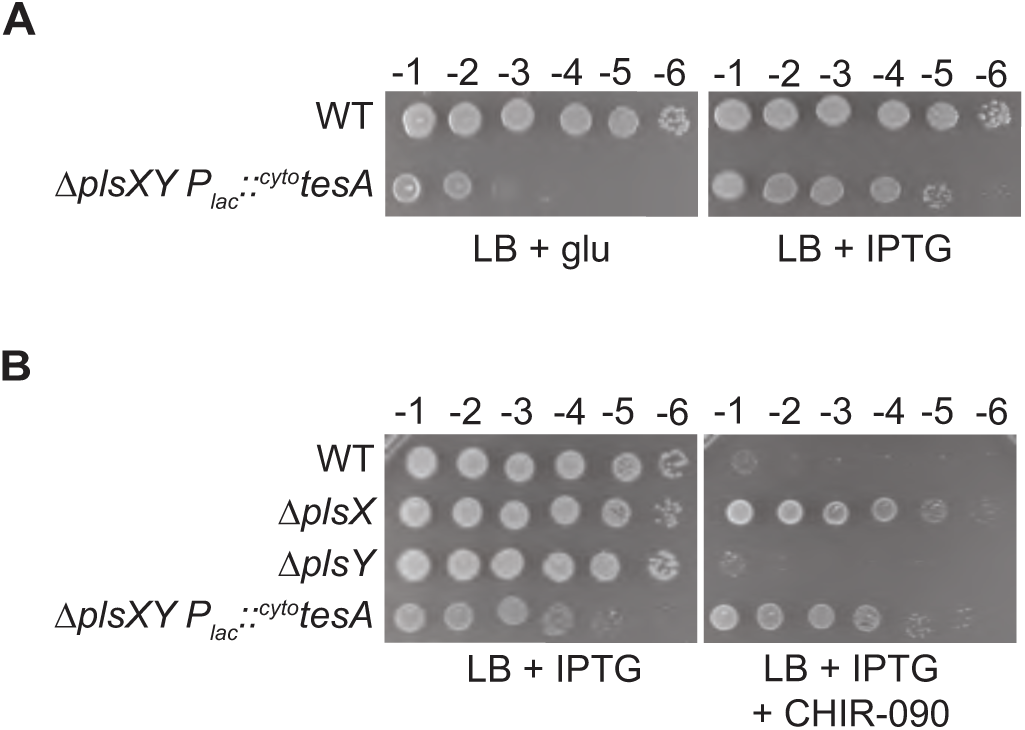
^cyto^TesA production rescues *ΔplsXY* synthetic lethality without influencing resistance to CHIR-090. **A.** Cells of the indicated strains were spotted on LB with 0.2 % glucose (left, glu) or LB with IPTG (right, 50 µM) and grown at 30°C for 17 hours before imaging. **B.** Cells of the indicated strains were spotted on LB with IPTG (1 mM) and CHIR-090 (0.075 µg/mL) as indicated. Plates were incubated at 30°C for 17 hours before imaging. In *ΔplsXY* cells ^cyto^TesA was produced from pTGD35 [*P_lac_:: ^cyto^tesA*].

**Figure S6:**
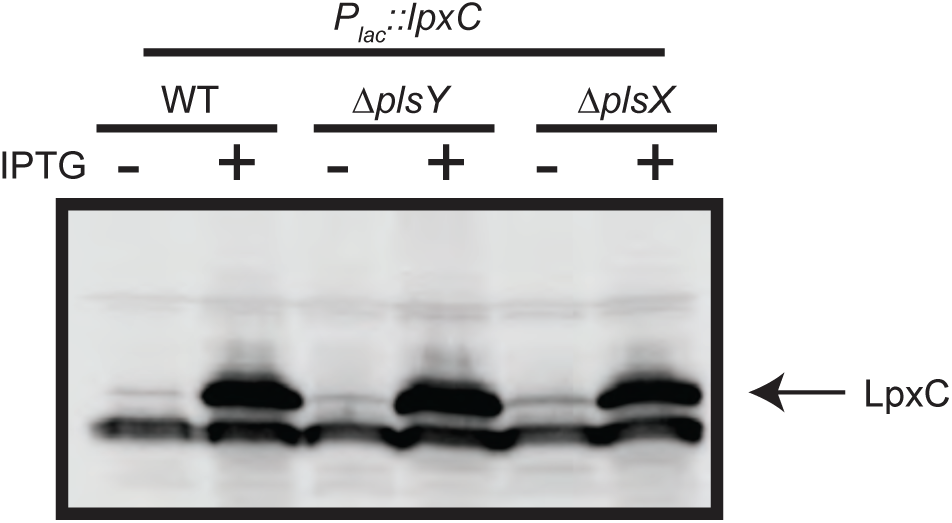
Quantification of LpxC overproduction. MG1655, *ΔplsX*, and *ΔplsY* containing pPR115 [*P_lac_::lpxC*] were grown with and without IPTG (1 mM) before extracts were prepared and LpxC was detected by immunoblotting with anti-LpxC antibodies. The band corresponding to LpxC is indicated by the arrow.

**Figure S7:**
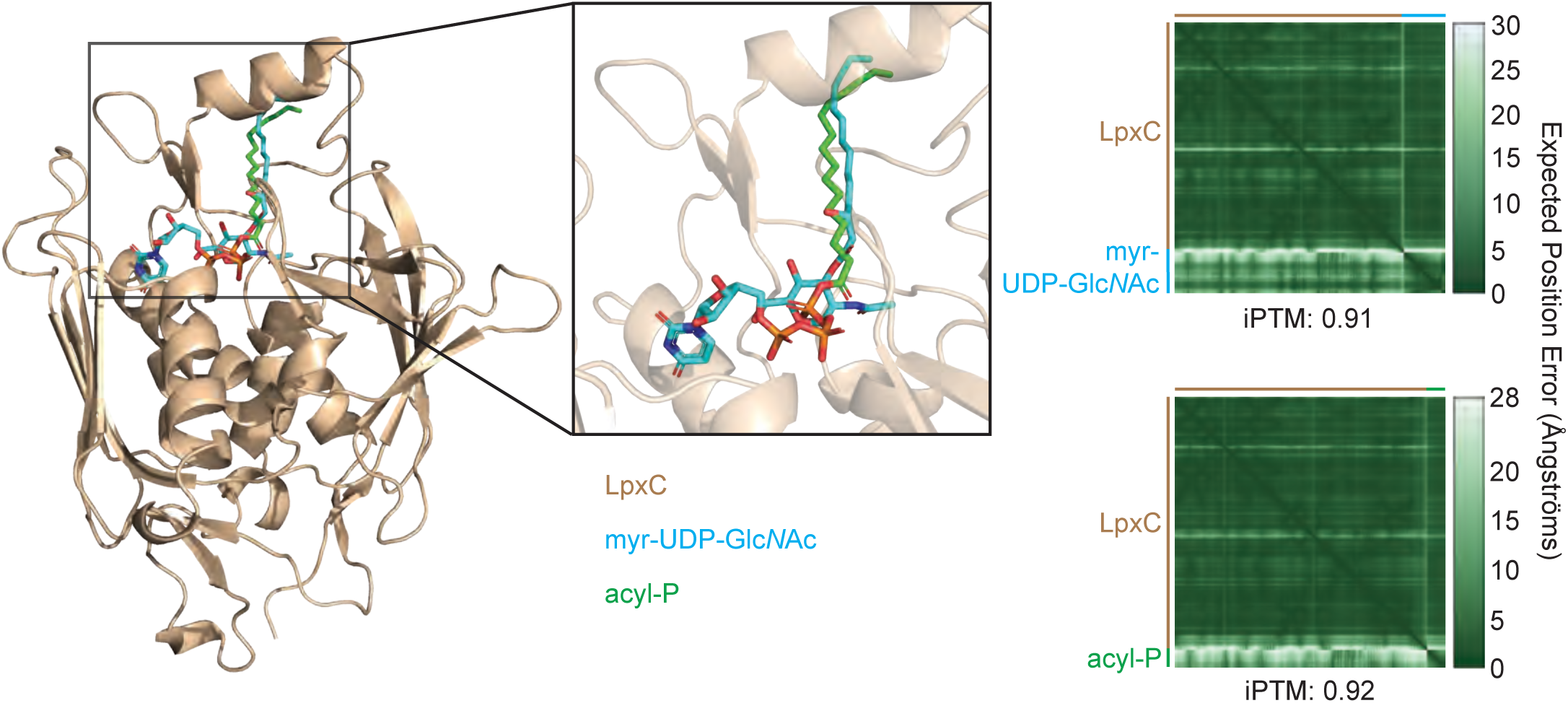
***E. coli* LpxC modeled with native substrate and acyl-P.** Predicted structure of E. coli LpxC (beige) interacting with its native substrate UDP-3-O-(R-3-hydroxymyristoyl)-N-acetylglucosamine (”myr-UDP-GlcNAc”, cyan) or palmitoyl-phosphate (”acyl-P”; green). The box shows a zoomed in view of the active site of LpxC. The prediction was modeled with AlphaFold 3. Predicted Alignment Error plots depicting the confidence of each molecule’s interaction with LpxC are shown to the right.

**Table S1:**
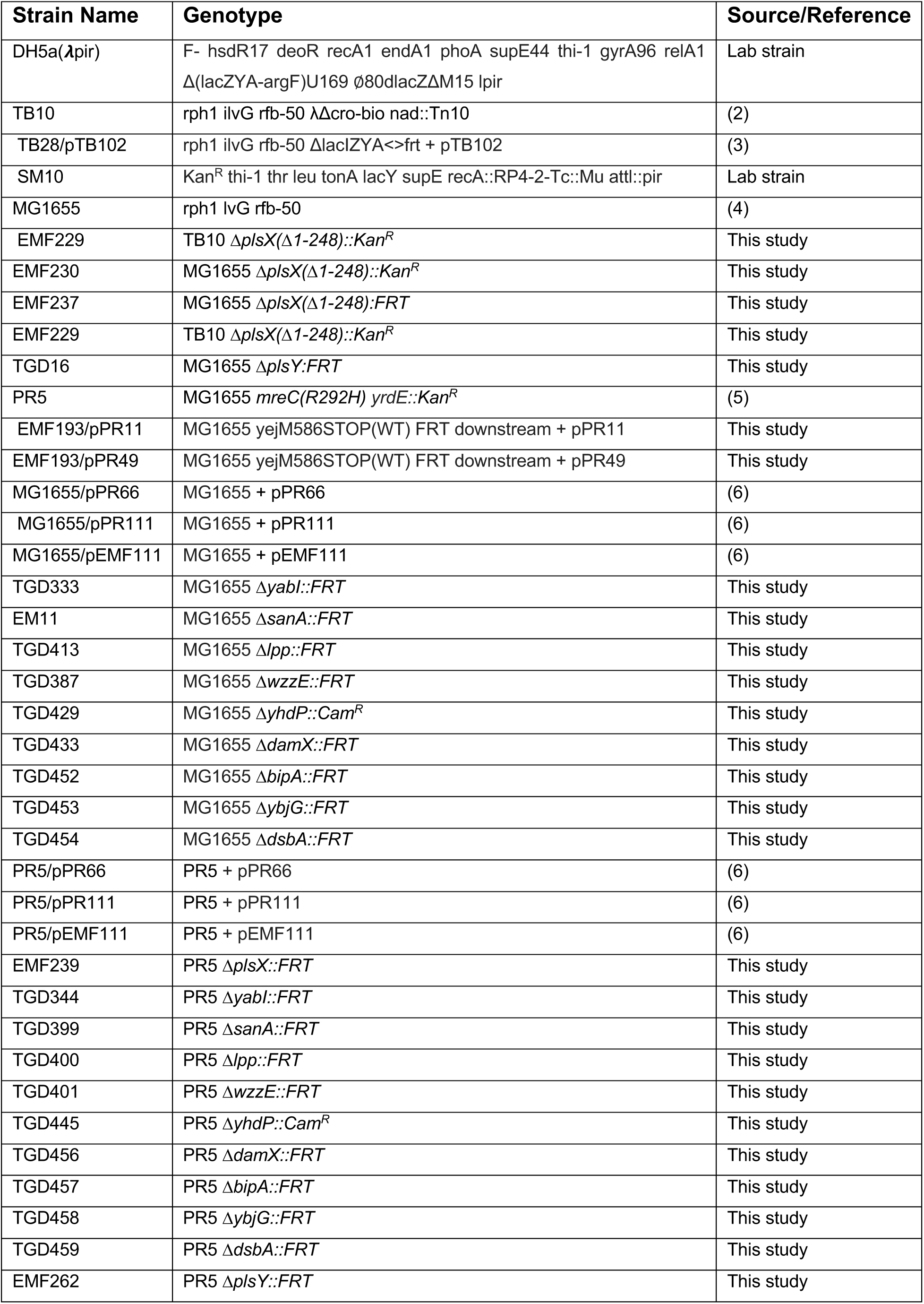
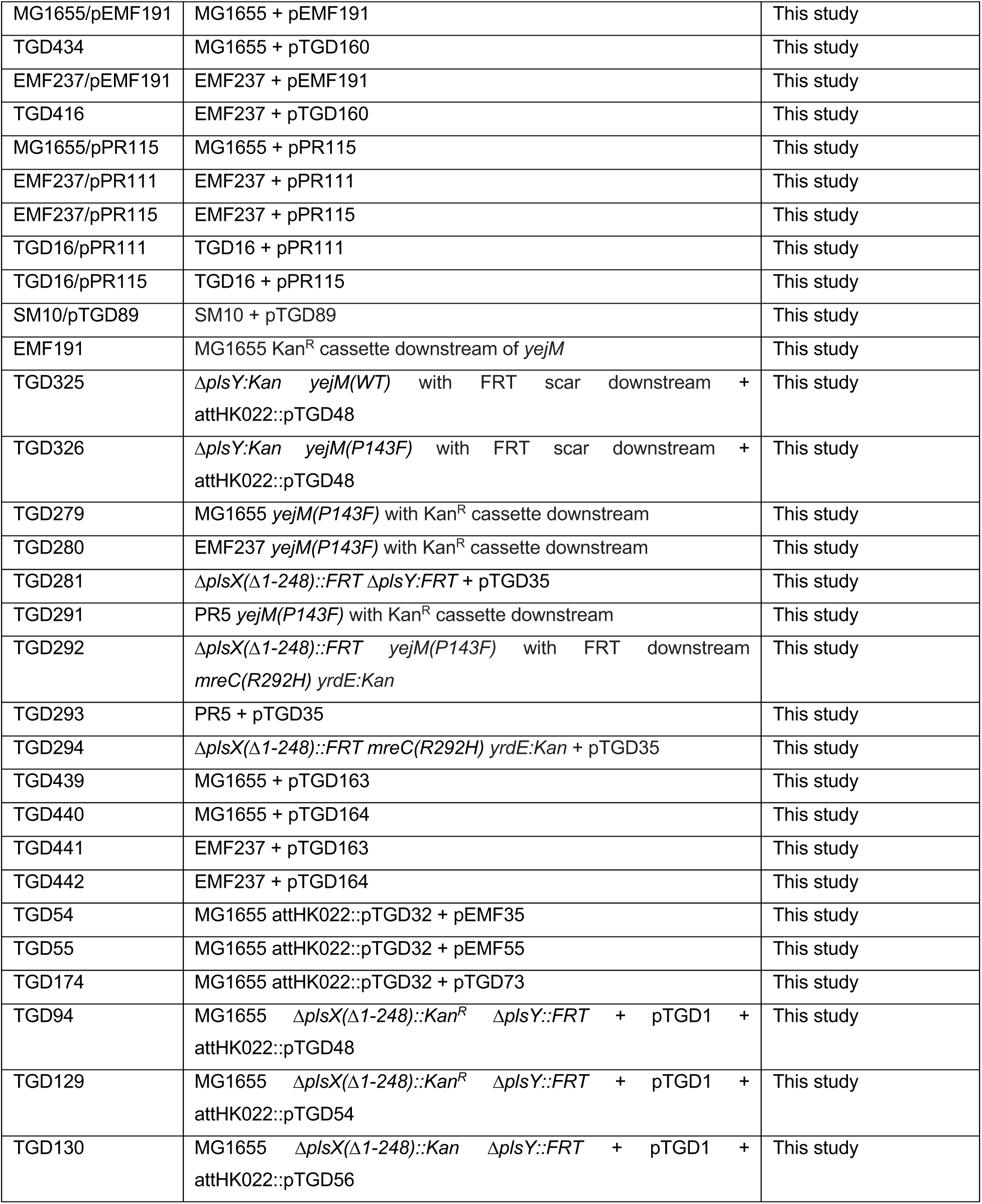
Strains used in this study.

**Table S2:**
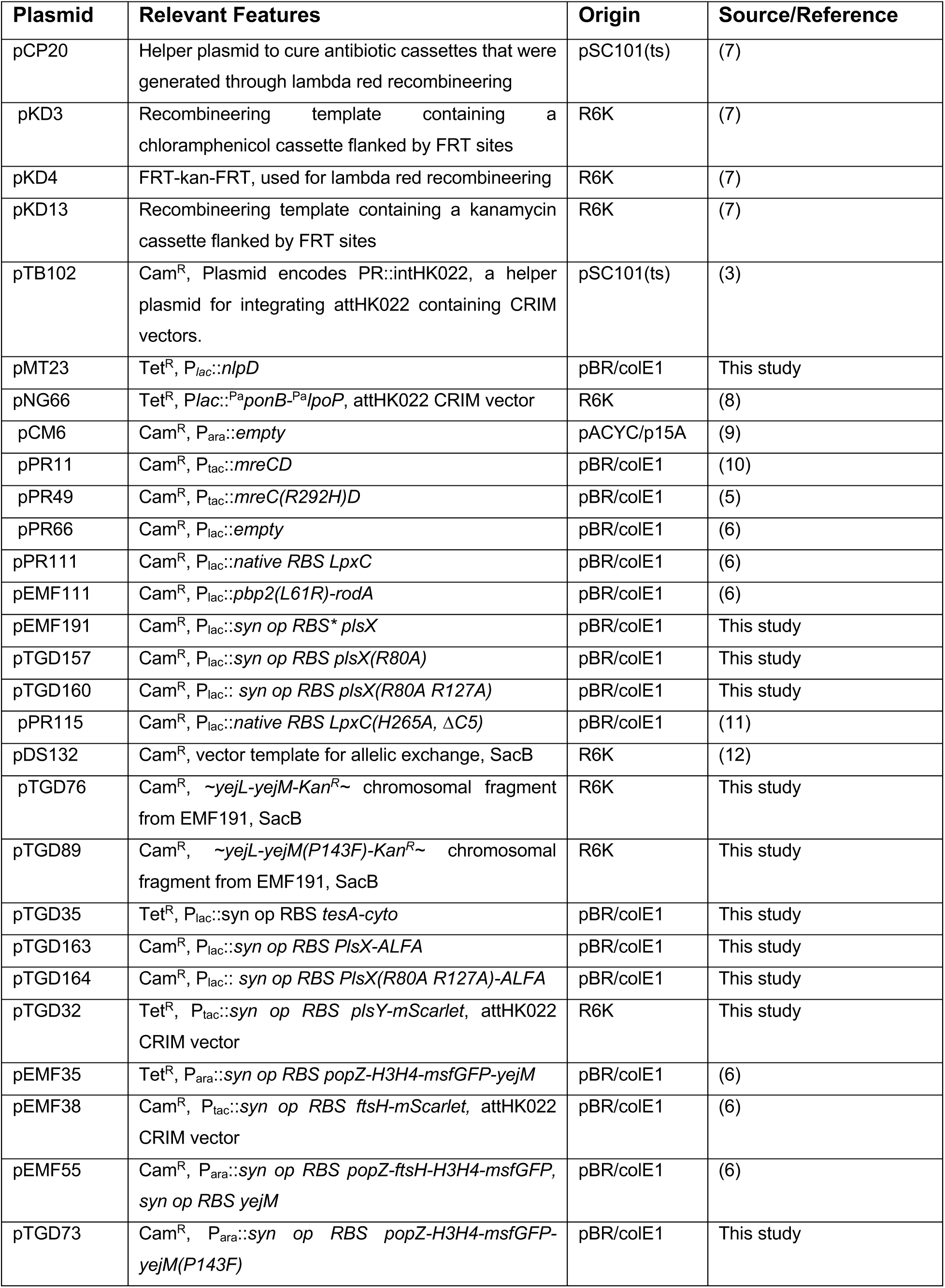
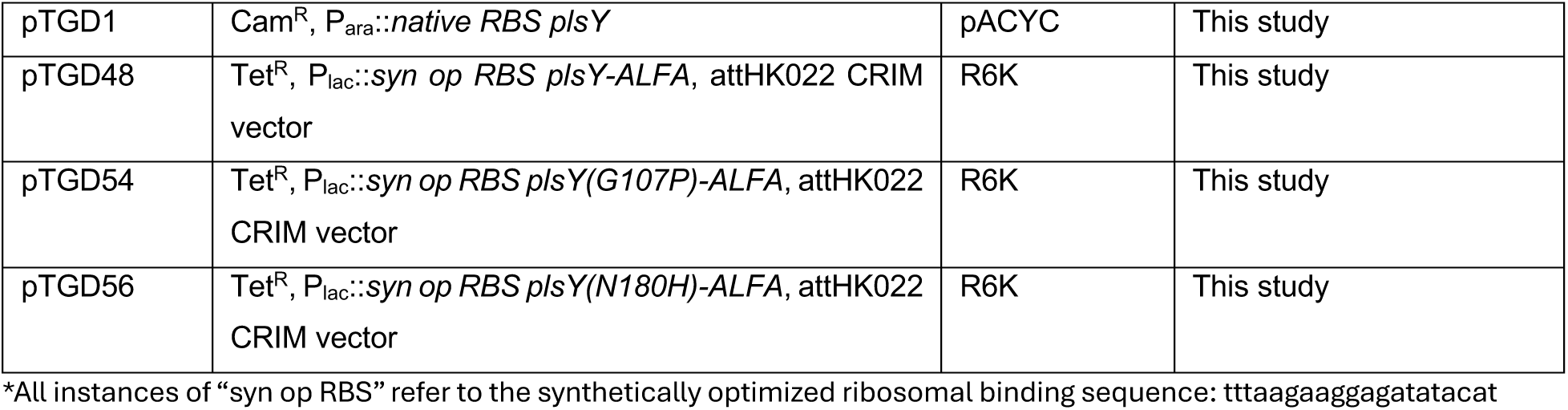
Plasmids used in this study.

**Table S3:**
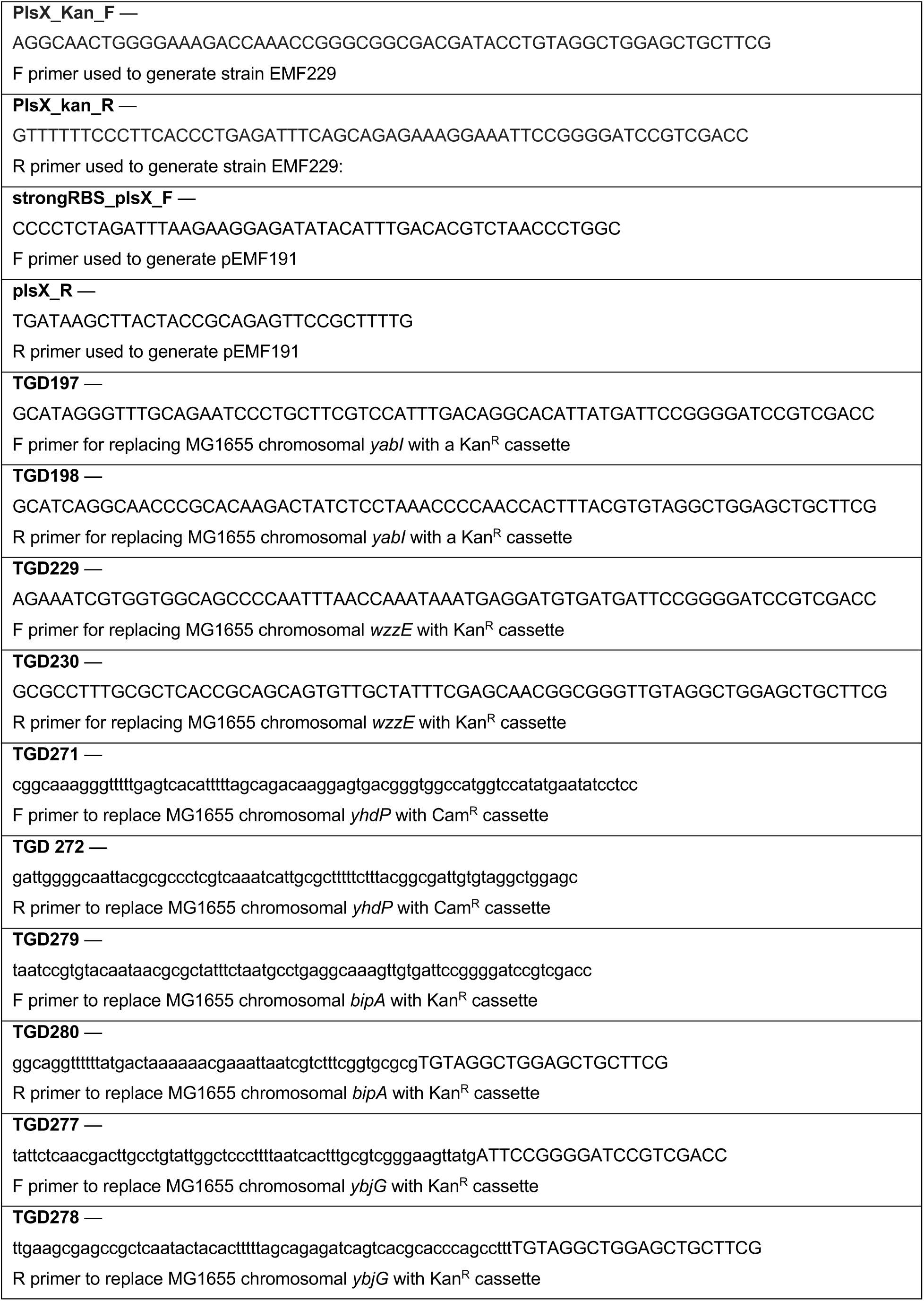
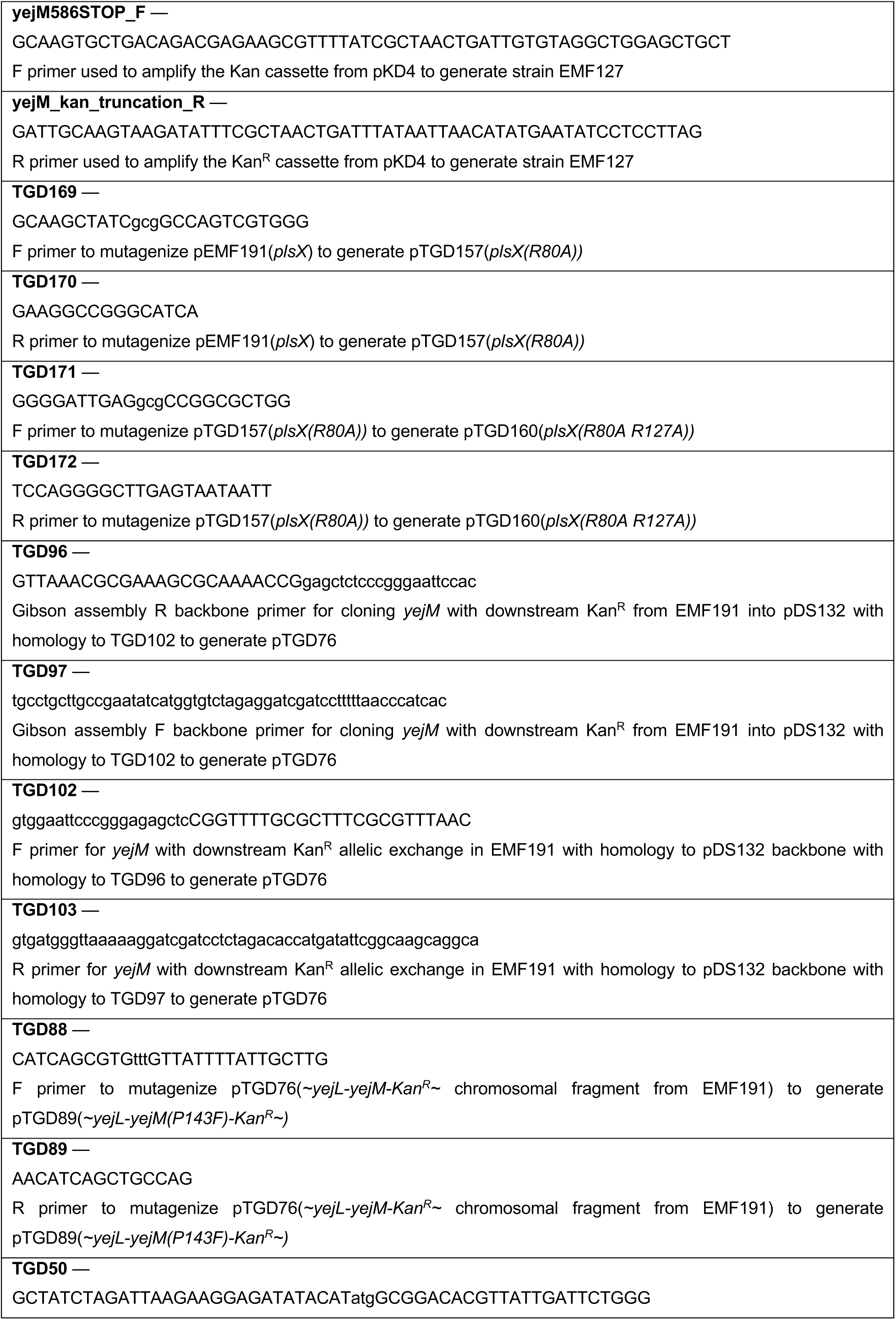
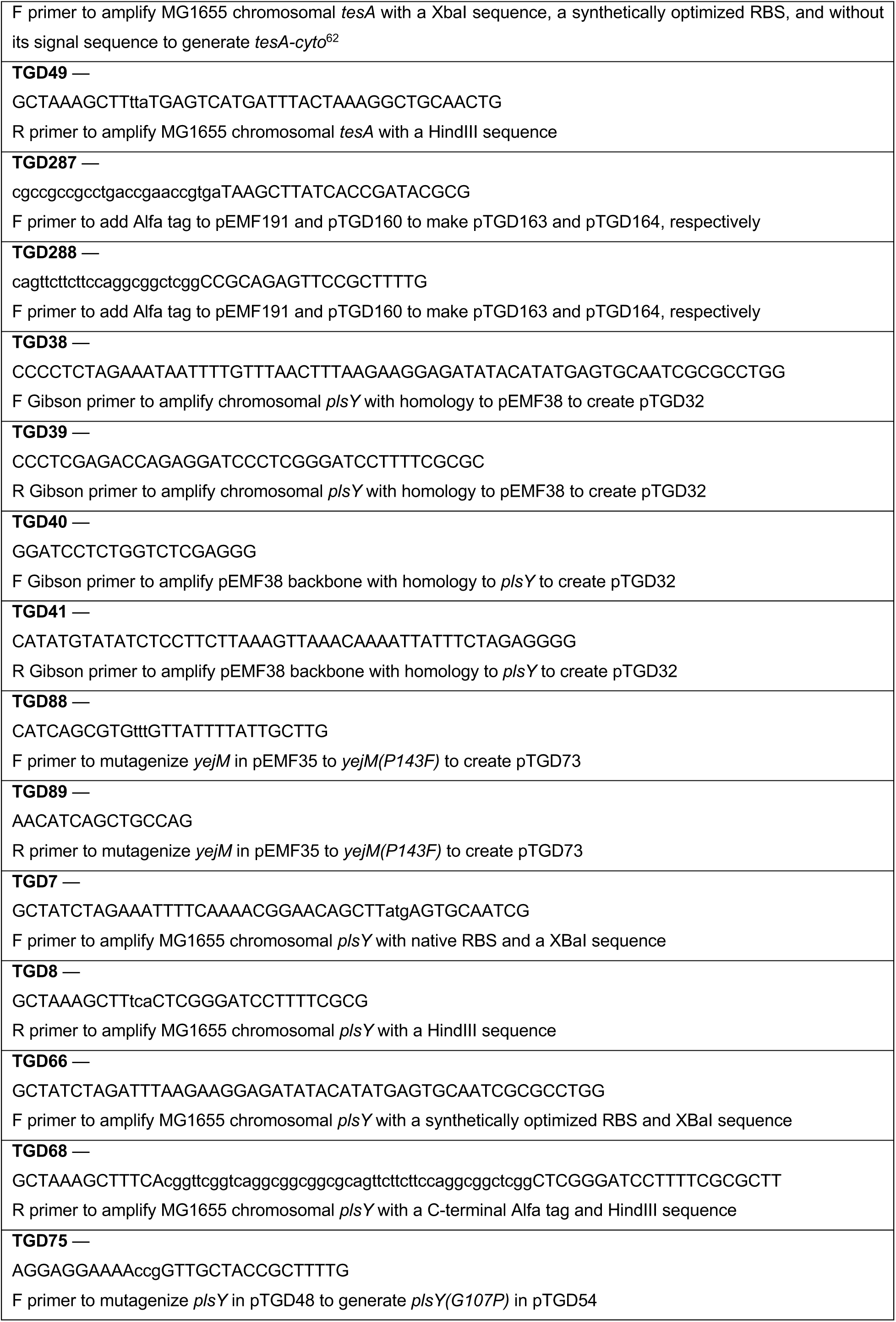
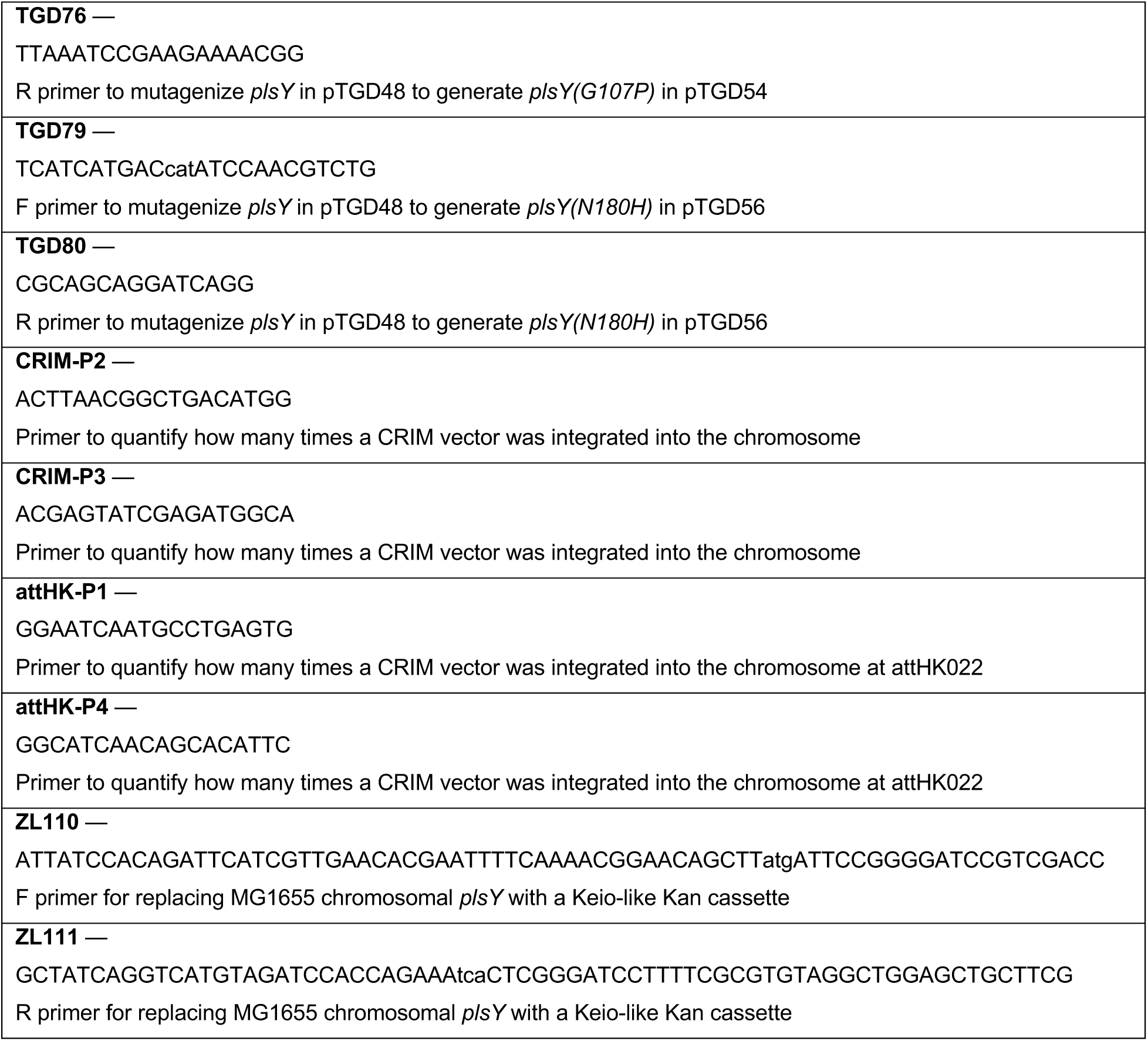
Primers used in this study.

**Table S4:**
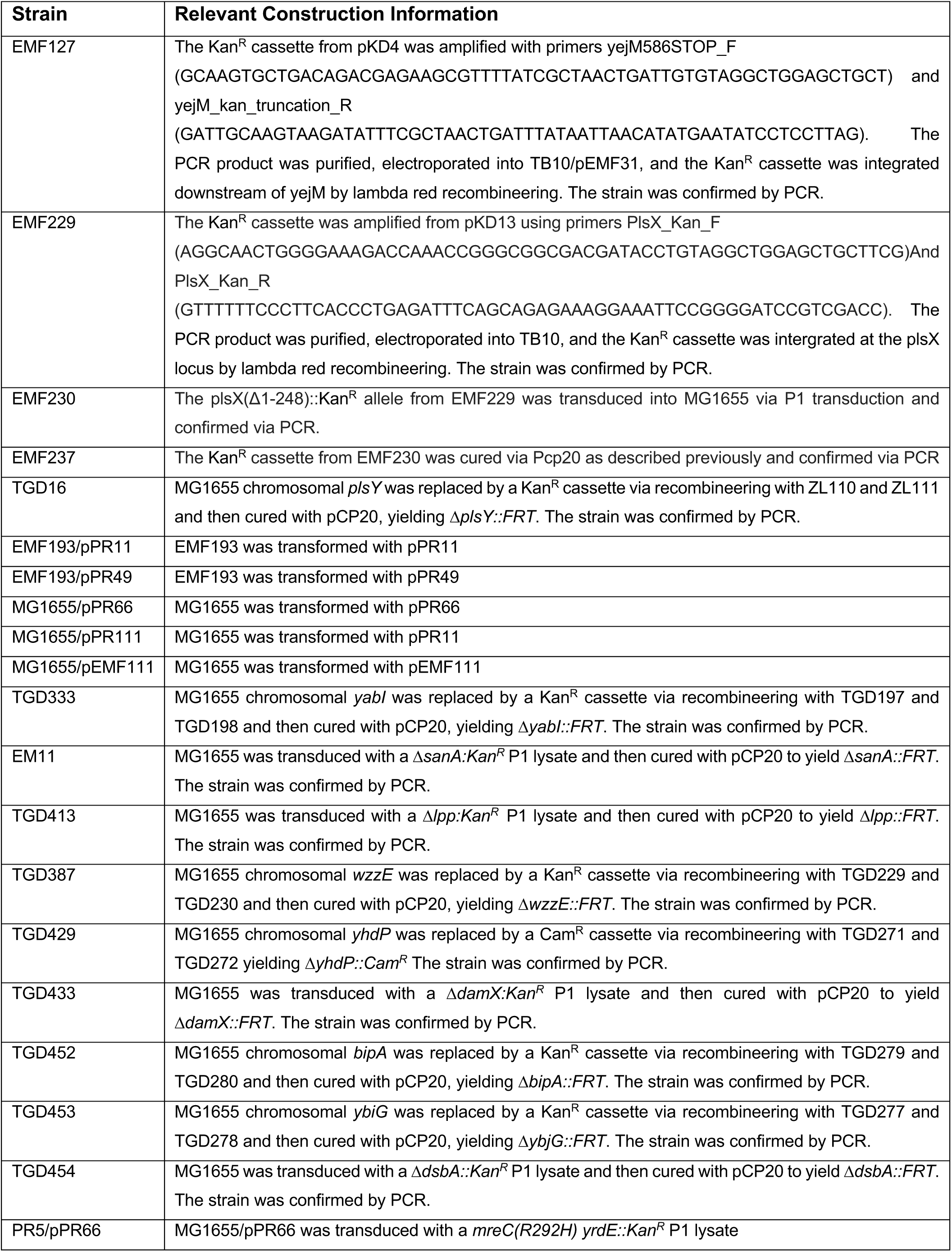
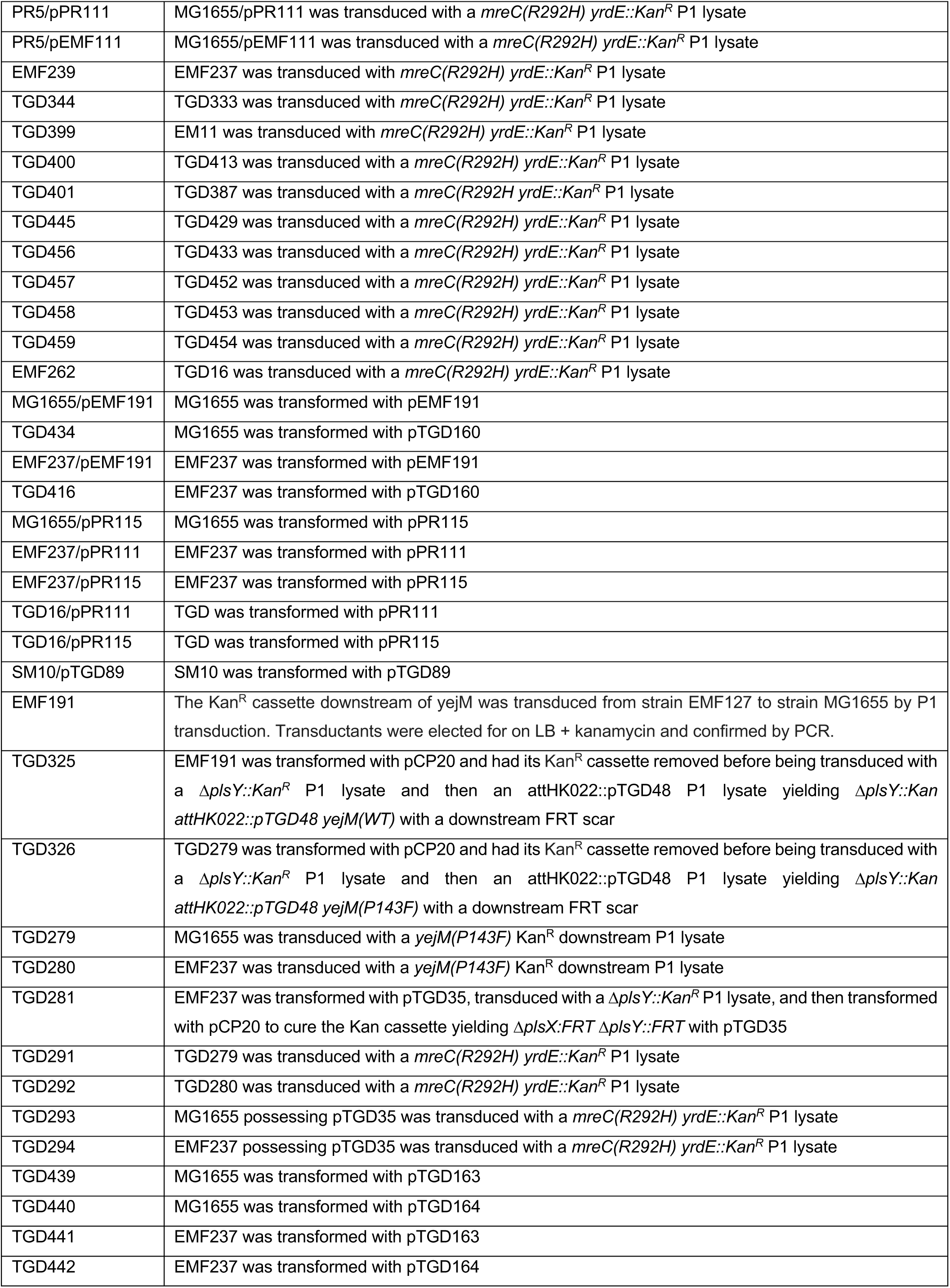
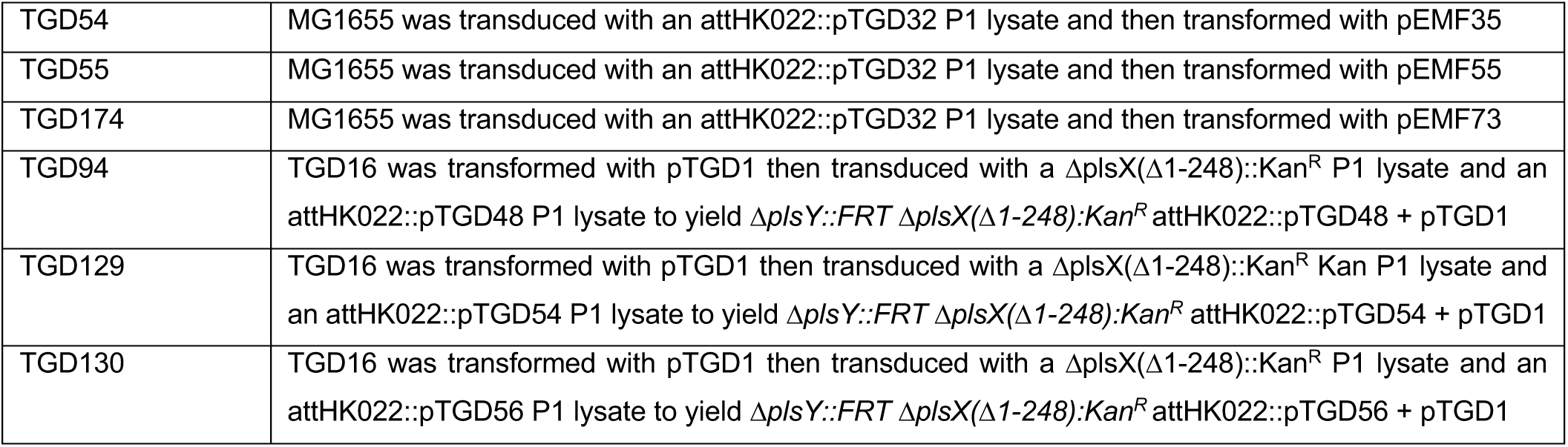
Strain construction information.

**Table S5:**
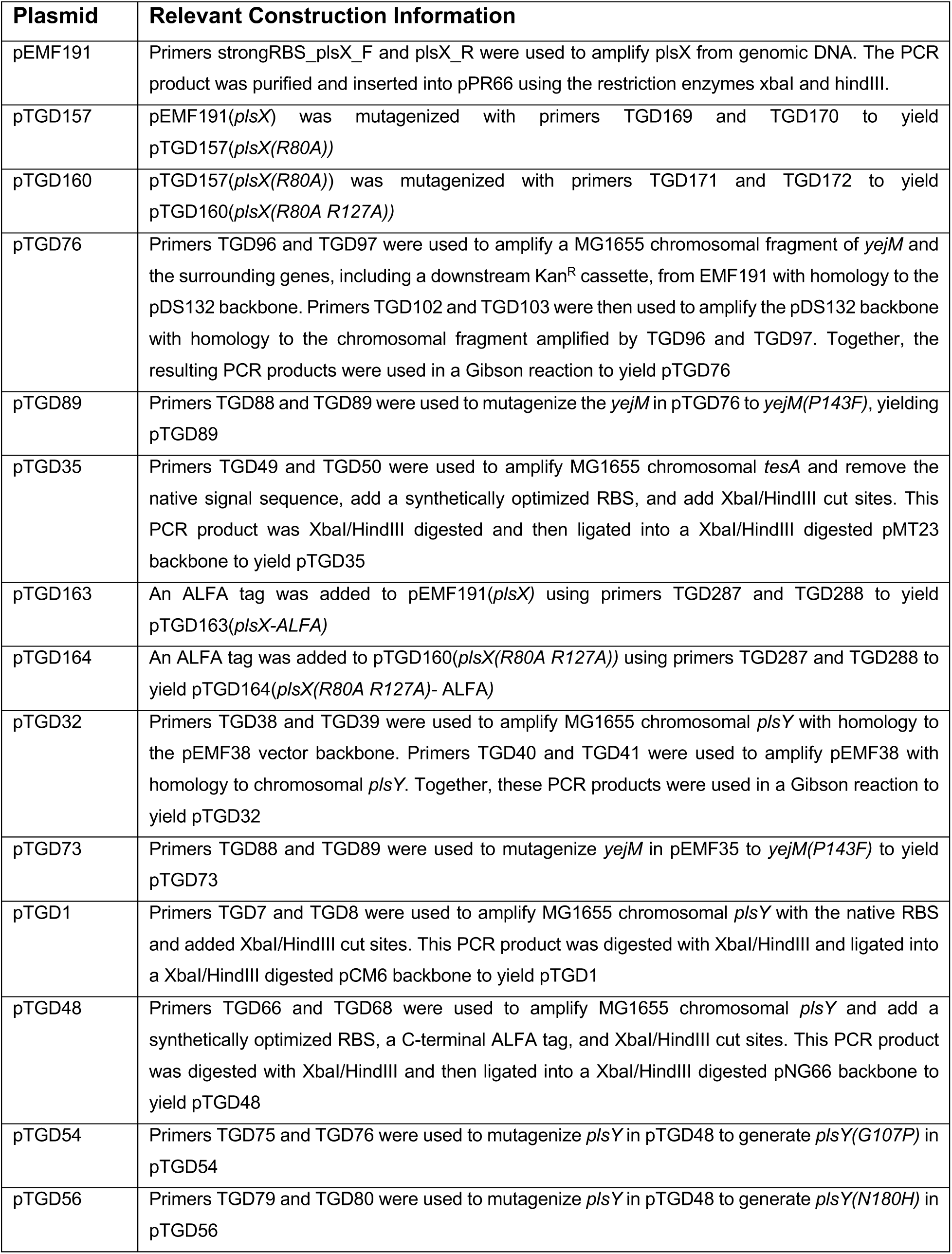
Plasmid construction information.

## Notes

### Competing Interest Statement

The authors have declared no competing interest.

